# Enolase inhibitors as therapeutic leads for *Naegleria fowleri* infection

**DOI:** 10.1101/2024.01.16.575558

**Authors:** Jillian E. Milanes, Victoria C. Yan, Cong-Dat Pham, Florian Muller, Samuel Kwain, Kerrick C. Rees, Brian N. Dominy, Daniel C. Whitehead, Steven W. Millward, Madison Bolejack, Jan Abendroth, Isabelle Q. Phan, Bart Staker, E. Ashley Moseman, Xiang Zhang, Xipeng Ma, Audriy Jebet, Xinmin Yin, James C. Morris

**Affiliations:** Eukaryotic Pathogens Innovation Center Department of Genetics and Biochemistry Clemson University, Clemson, SC 29634; Department of Cancer Systems Imaging UT MD Anderson Cancer Center, Houston TX 77030; Sporos Bioventures, 3000 Bissonnet, Belmont Suite 5303, Houston, TX 77005; Eukaryotic Pathogens Innovation Center Department of Chemistry, Clemson University, Clemson, SC 29634; UCB BioSciences, Bainbridge, Island, WA, 98110; Seattle Structural Genomics Center for Infectious Disease Center for Global Infection Disease Research, Seattle Children’s Research Institute, Seattle, WA 98109; Department of Immunology, Duke University School of Medicine, Durham, NC 27710; Department of Chemistry, University of Louisville, Louisville, KY 40208

**Keywords:** enolase, free-living amoebae, glycolysis, inhibitors, *Naegleria fowleri*

## Abstract

Infections with the pathogenic free-living amoebae *Naegleria fowleri* can lead to life-threatening illnesses including catastrophic primary amebic meningoencephalitis (PAM). Efficacious treatment options for these infections are lacking and the mortality rate remains >95% in the US. Glycolysis is very important for the infectious trophozoite lifecycle stage and inhibitors of glucose metabolism have been found to be toxic to the pathogen. Recently, human enolase 2 (ENO2) phosphonate inhibitors have been developed as lead agents to treat glioblastoma multiforme (GBM). These compounds, which cure GBM in a rodent model, are well-tolerated in mammals because enolase 1 (ENO1) is the predominant isoform used systemically. Here, we describe findings that demonstrate that these agents are potent inhibitors of *N. fowleri* ENO (*Nf*ENO) and are lethal to amoebae. In particular, (1-hydroxy-2-oxopiperidin-3-yl) phosphonic acid (HEX) was a potent enzyme inhibitor (IC_50_ value of 0.14 ± 0.04 µM) that was toxic to trophozoites (EC_50_ value of 0.21 ± 0.02 µM) while the reported CC_50_ was >300 µM. Molecular docking simulation revealed that HEX binds strongly to the active site of *Nf*ENO with a binding affinity of –8.6 kcal/mol. Metabolomic studies of parasites treated with HEX revealed a 4.5 to 78-fold accumulation of glycolytic intermediates upstream of *Nf*ENO. Last, nasal instillation of HEX increased longevity of amoebae-infected rodents. Two days after infection, animals were treated for 10 days with 3 mg/kg HEX, followed by one week of observation. At the conclusion of the experiment, eight of 12 HEX-treated animals remained alive (resulting in an indeterminable median survival time) while one of 12 vehicle-treated rodents remained, yielding a median survival time of 10.9 days. Brains of six of the eight survivors were positive for amoebae, suggesting the agent at the tested dose suppressed, but did not eliminate, infection. These findings suggest that HEX is a promising lead for the treatment of PAM.

## INTRODUCTION

Human infections by the pathogenic free-living amoebae (pFLA) can lead to life-threatening illnesses, a situation exacerbated by the lack of useful therapeutic drugs for successful interventions. Perhaps the best-known pFLA, *Naegleria fowleri* establishes infections after trophozoites in contaminated water are introduced into the nasal passages and ultimately penetrate the cribriform plate to infect the brain. This infection, known as primary amebic meningoencephalitis, is almost invariably fatal due to tissue damage caused by the parasite and concurrent hemorrhagic necrosis of the brain leading to lethal cerebral edema or multiorgan system failure (1). Due to the high mortality rate, poor therapeutic options, and lack of awareness and detection, this organism is classified as a category B NIAID emerging infectious pathogen.

We have previously found that glucose metabolism is critical to the viability of the infectious trophozoite lifecycle stage (2). This pathway is anticipated to impinge on multiple critical cellular functions, as it provides the cell with ATP and key metabolic intermediates. For example, glucose 6-phosphate (G6P) generated by the first step in the process is needed for the pentose phosphate pathway (PPP), which generates essential metabolic products including reducing equivalents and sugar nucleotide biosynthetic intermediates.

The role of glucose metabolism during infection is unclear. A comparison of transcriptomes from continuously cultured *N. fowleri* to amoebae recently isolated from an infected brain revealed that multiple metabolic genes were upregulated including those involved in fatty acid metabolism as well as glycolysis, further clouding the role of glucose metabolism during infection (3). Growth of *N. gruberi,* a non-pathogenic *Naegleria*, requires glucose, even though the organism preferentially metabolizes fatty acids (4). This suggested that some level of glycolysis or gluconeogenesis is likely also essential for *N. fowleri* viability, a finding supported by the lack of trophozoite growth when cultured in the absence of glucose (2).

Lin and colleagues have recently described the development of phosphonate inhibitors of human ENO2, the enzyme that interconverts 2-phosphoglycerate (2-PG) to phosphoenolpyruvate (PEP) in glycolysis and gluconeogenesis, for the treatment of *ENO1*^−/−^ GBM (5). These inhibitors selectively killed *ENO1*-deleted glioma cells and eliminated intracranial orthotopic *ENO1*-deficient tumors in mice. One of the lead compounds, HEX, had an EC_50_ value of 1.3 μM against *ENO1*^−/−^ glioma cells but was very well-tolerated by *ENO1*-rescued glioma and other control cells, with CC_50_ values >300 μM. HEX also almost completely inhibited intracranial tumor growth and these impacts did not require a breached blood-brain barrier. Importantly, therapeutic doses were at levels well-tolerated in non-human primates (NHPs), with IP delivery of HEX tolerated to 300 mg/kg (body weight) without adverse effects, yielding plasma concentrations of about 750 μM for several hours with a plasma elimination half-life in non-human primates of 61 minutes.

Given the importance of glycolysis to *N. fowleri* and the known distribution of the phosphonate ENO2 inhibitors to the brain, we have assessed the activity of these compounds as potential anti-amebic agents. This includes scoring the impact of the inhibitors against the *N. fowleri* ENO (*Nf*ENO) through a series of biochemical assays and against amoebae trophozoites, both *in vitro* and in a rodent model of disease. The findings of these efforts suggest the agents could be a useful component for treatment of *N. fowleri* infections.

## MATERIALS AND METHODS

### Chemicals and Reagents

β-nicotinamide adenine dinucleotide phosphate (NADP^+^), adenosine diphosphate (ADP), and dimethyl sulfoxide (DMSO) were obtained from Fisher Scientific (Pittsburgh, PA, USA). Pyruvate kinase/lactate dehydrogenase (PK/LDH) from rabbit muscle and 2-phophoglyceric acid disodium salt hydrate (2-PG) were both purchased from MilliporeSigma (Burlington, MA, USA). Human ENO1, 2, and 3 were graciously provided by Dr. Paul Leonard (MD Anderson Cancer Center Core for Biomolecular Structure and Function).

The compounds (1-hydroxy-2-oxopyrrolidin-3-yl) phosphonic acid (deoxy-SF2312), (1-hydroxy-2-oxopiperidin-3-yl) phosphonic acid (HEX), and (1-hydroxy-2-oxoazepan-3-yl) phosphonic acid (HEPTA), were synthesized following published procedures reported in the literature (5, 6). Please see the synthetic scheme (Fig. S1) and spectroscopic characterization (Fig. S2).

### Protein purification, crystallization, and structure determination

Protein purification, crystallization, and structure determination were conducted by the Seattle Structural Genomics Center for Infectious Disease (SSGCID) (7, 8) following standard protocols described previously (9–11). (See Supplemental Materials for further information.) Briefly, *E.coli* codon optimized sequence encoding residues 2-512 of the full length protein (NF0118810, AmoebaDB, https://amoebadb.org/amoeba/app/) was used to develop five constructs composed of residues 14-512, 28-512, 44-512, 62-512, 80-512 that were predicted by Xtalpred (https://xtalpred.godziklab.org/XtalPred-cgi/xtal.pl) to be optimal for crystallization. Following cloning into pBG1861 (10), which appends a non-cleavable 6xHis N-terminal tag to fusion proteins, constructs were transformed into *E. coli* BL21(DE3)R3 Rosetta cells. Two constructs, *Nf*ENO(44-512) and *Nf*ENO(62-512), yielded soluble protein. Please note that an alternate N-terminal translation start site has been annotated in Uniprot as A0A6A5BXC3, yielding a predicted 485 residue polypeptide that includes residues equivalent to 28-512 of NF0118810 (see Fig. S3 and Supplemental Table S1).

Following large scale growth in auto-induction media (12), bacteria were lysed by sonication and clarified lysate was subject to sequential Ni^2+^-affinity and size-exclusion chromatography (SEC). The SEC peak fractions eluted as a single large peak consistent with a molecular mass of about 50 kDa, suggesting a monomeric enzyme, which was consistent with the size of the denatured purified protein as determined by SDS-PAGE. Peak fractions were pooled and concentrated.

Purified *Nf*ENO(44-512) and *Nf*ENO(62-512) were screened for crystallization in 96-well sitting-drop plates against the JCSG++ HTS (Jena Bioscience) and PACT premier HT96 (Molecular Dimensions) crystal screens. In addition to protein-only crystallizations, co-crystallization trials with 2-PG (5 mM) and HEX (5 mM) were also pursued. Only *Nf*ENO(44-512) in complex with 2-PG crystallized to form diffraction quality crystals, which were flash-frozen by plunging directly into liquid nitrogen without cryoprotectant exchange.

X-ray data were collected at 100 K on beamline 21-ID-F, LS-CAT, at the Advanced Photon Source, Argonne National Laboratory. Data were processed with XDS reduced with XSCALE (13) and the structure was solved by molecular replacement using the structure of *B. subtilus* ENO (PDB 4A3R) as a search model (14). Structures were refined using iterative cycles of Phenix (15) followed by manual rebuilding of the structure using Coot (16) to yield a resolution of 1.95 Å, with the quality of all structures checked using MolProbity (17). All data-reduction and refinement statistics are shown in Supplemental Tables S2 and S3. Structure figures were prepared and analyzed using PyMOL (v.1.5; Schrodinger) and coordinates and structure factors have been deposited in the Protein Data Bank www.rcsb.org (18) with accession number 7UGH.

### ENO Assays

ENO was assayed in triplicate using a coupled reaction to measure enzyme activity (19). Briefly, enzyme (50 nM) in assay buffer (100 mM HEPES, pH 8.0, 3.3 mM MgSO_4_) was loaded in black 96-well plates with or without inhibitors. To start the reaction, substrate buffer (3.75 mM 2-PG, 1.75 mM ADP, 1 U of pyruvate kinase/lactate dehydrogenase, and 0.4 mM NADH in assay buffer) was added to each well. The rate of reduction in fluorescence (excitation: 360 nm; emission: 460 nm), a measure of NAD production, was then scored on a Biotek Synergy H1 microplate reader every 20 seconds for 4 minutes. Kinetic analyses were performed with Prism 9.0 (GraphPad Software, San Diego, CA) using the Michaelis-Menten model. For assays involving inhibitors, compounds (in DMSO) were incubated at RT with enzyme in assay buffer (pH 7) for 15 minutes prior to initiation of the reaction by addition of the substrate. All assays were performed in triplicate with vehicle control (DMSO) included.

### Ligand preparation, optimization, and molecular docking

The chemical structures of HEX, (1,5-dihydroxy-2-oxopyrrolidin-3-yl) phosphonic acid (SF2312), deoxy-SF2312, and HEPTA were drawn with ChemOffice professional 19 suite (PerkinElmer, Waltham, MA), and three-dimensional (3D) structures were generated with VeraChem Vconf (VerChem LLC, Germantown, MD). The 3D structures were optimized by Gaussian 16 suite (Gaussian Inc., Wallingford, CT) with Density Functional Theory (DFT), employing the B3LYP/6-311G (d,p) level of theory (20). The phosphonate-bearing stereocenter was modeled with the (R)-configuration by analogy to the enantiomer of HEX that crystallized in human ENO (PDB 5IDZ). Since SF2312 likely exists as a mixture of *cis* and *trans* diastereomers at physiological pH, we modeled both diastereomers independently while maintaining the phosphonate-bearing stereocenter in the (R)-configuration (5, 6, 21). The *Nf*ENO 3D crystal structure (PDB 7UGH) was retrieved from the RCSB protein data bank. The protein was then prepared for the docking analysis by first removing the co-crystallized ligand, heteroatoms, and water molecules using Pymol Molecular Graphics 2.0 (Schrödinger LLC, New York, NY). The optimized ligands and the protein were further organized using AutoDock Tools (The Scripps Research Institute, La Jolla, CA) to convert all structures into pdbqt formats. The grid box was placed around the region of the active site of the protein. The size of the grid box was kept at 48, 52, and 52 for X, Y, and Z respectively with the center of the grid box maintained at –18.910, –1.626, and –15.343 respectively for X, Y, and Z. The molecular docking studies were carried out *in vacuo* with AutoDock vina using specific docking parameters and scoring functions (22). The binding affinities of ligands were measured in kcal/mol as a unit for a negative score (22). The binding conformation with the highest negative value was taken as the best pose for the corresponding protein-ligand complex. Subsequently, the best binding pose of each complex was analyzed using Pymol and Discovery Studio (Dassault Systèmes, Waltham, MA) to reveal the protein-ligand interactions.

### *In vitro* growth inhibition and selection for resistance

*N. fowleri* strain Nf69 (ATCC 30215) trophozoites and *N. gruberi* strain NEG (ATCC 30223) were cultured axenically as previously described (2). Briefly, parasites were seeded at 1 × 10^4^ cells/mL (*N. fowleri*) or 5 × 10^3^ cells/mL (*N. gruberi*) in 100 µL of the indicated medium in white TC-treated 96-well plates (Thermo Scientific Nunc Microwell 136101). For assays under standard growth conditions (“+ glc*”), N. fowleri* cells were seeded into Nelson’s Complete Media (NCM, 0.17% liver infusion broth (BD Difco, Franklin Lakes, NJ), 0.17% glucose, 0.012% sodium chloride, 0.0136% potassium phosphate monobasic, 0.0142% sodium phosphate dibasic, 0.0004% calcium chloride, 0.0002% magnesium sulfate, 10% heat-inactivated fetal bovine serum (FBS), 1% penicillin-streptomycin) at 37 °C. For assays in modified medium (“-glc/+gly”), cells were seeded into the media without glucose but supplemented with 10 mM glycerol. For standard growth of *N. gruberi*, cells were seeded into PYNFH-complete media (1.0% peptone, 1.0% yeast extract, 0.1% yeast nucleic acid, 0.0015% folic acid, 0.0001% hemin, 0.0362% potassium phosphate monobasic, 0.05% disodium phosphate, 10% heat-inactivated fetal bovine serum (FBS), 1% penicillin-streptomycin) at 25 °C. After culturing for 48 hr (37 °C, 5% CO_2_ for *N. fowleri*; 25 °C, 5% CO_2_ for *N. gruberi*) in the presence of compound or vehicle, plates were equilibrated to RT for 15 min and CellTiter Glo reagent (Promega, Madison, WI, USA) was added followed by orbital shaking for 2 min. After an additional incubation (RT, 10 min), luminescence was scored on a BioTek Synergy H1 mircoplate reader. Parasite density was scored and kinetic analyses were performed with Prism 9.0 (GraphPad Software, San Diego, CA) using the Michaelis-Menten model.

To generate resistance in *N. fowleri*, cells were seeded in 75 cm^2^ TC-treated flasks (Falcon 353135) at 1 × 10^4^ cells/mL in the presence of HEX, starting at the calculated (using GraphPad software) EC_25_ concentration. After one week, HEX levels were increased to the EC_50_ concentration for an additional two weeks. At this point, the compound was increased weekly (2 × EC_50_ value, 5 × EC_50_ value, and 10 × EC_50_ value), with cells passaged as needed throughout in the presence of the compound. Amoebae were then cultured for one month in 10 × EC_50_ levels of HEX before performing viability assays.

### Analysis of metabolites after HEX treatment

For unbiased metabolomics analyses, *N. fowleri* trophozoites were grown to near 100% confluency in 6-well TC-treated plates (Thermo Scientific Nunclon Delta Surface 140685). HEX (25 µM) or vehicle control was added to appropriate wells for an additional 6 hours. Samples were then washed three times in RT PBS prior to the addition of ice-cold CH_3_CN. Plates were then incubated at – 20 °C for 20 min, followed by the addition of ice-cold nanopure H_2_O and removal of the solution to sterile microcentrifuge tubes. This process was repeated twice, and the total volume of extraction material was adjusted to 2.5 × 10^6^ cell equivalents per sample. Tubes were stored at –80°C prior to analysis.

### Metabolomic analysis

Each cell sample was vortexed with glass beads for 5 min in a MM200 Mix Grinder (Retsch USA, Newtown, PA, USA). The mixture was centrifuged at 14,000 rpm for 20 min at 4 °C. The supernatant was transferred into a new tube and lyophilized overnight. The dried sample was reconstituted in 60 µL 50% CH_3_CN and analyzed in random order on a parallel two-dimensional liquid chromatography-mass spectrometry (2DLC-MS) system, a Thermo DIONEX UltiMate 3000 HPLC system (Thermo Fisher Scientific, Waltham, MA, USA) coupled with a Thermo Q Exactive HF Hybrid Quadrupole-Orbitrap Mass Spectrometer (Thermo Fisher Scientific, Waltham, MA, USA). The UltiMate 3000 HPLC system was equipped with a hydrophilic interaction chromatography (HILIC) column and a reversed-phase chromatography (RPC) column configured in parallel mode. Each sample was analyzed by the 2DLC-MS under both positive (+) and negative (–) modes to obtain full MS data for metabolite quantification. A pooled sample was also prepared for each sample group and analyzed by 2DLC-MS/MS to acquire MS/MS spectra at three collision energies (20, 40, and 60 eV) for metabolite identification. Detailed 2DLC-MS operation parameters and data analysis methods were reported in our previous publications (23–25).

### Rodent model of infection

Twenty-four CD rats (12 females, 12 males, aged 42-48 days, Charles River) housed in pairs were infected by nasal instillation with 2.5 × 10^5^ *N. fowleri* strain 69 trophozoites that had been isolated after passage through mice to ensure virulence. Rats were provided unlimited food and water throughout the duration of the experiment. The two groups consisted of equal numbers of males and females. Groups received either vehicle (PBS) or HEX (3 mg/kg) delivered by nasal instillation (25 µL/nare) for 10 consecutive days starting 48hr after infection. Rodents were monitored carefully and euthanized when symptoms of infection, including failure to respond to stimuli or seizures, were evident. After 17 days, surviving animals were sacrificed and brains were removed to assess cure. Rats were considered cured if amoebae from isolated brains failed to grow in culture after two weeks.

### Ethics Statement

Rats were purchased from Charles River Laboratories and maintained in the Association for Assessment and Accreditation of Laboratory Animal Care (AAALAC) accredited, PHS assured, USDA registered Godley Snell Research Center at Clemson University. All protocols used in this study were approved by the Clemson University Institutional Animal Care and Use Committee (CU PHS Assurance Number D16-00435 (A3737-01) (Protocol #2023-0121) and performed in accordance with AAALAC International standards.

## RESULTS

*N. fowleri* trophozoites are reliant on glycolysis for normal growth *in vitro*, suggesting that inhibitors of enzymes in the pathway could serve as useful anti-amebic compounds. Supporting this supposition, inhibitors of glucokinase, the first enzyme in the pathway, have been identified that have modest amoebicidal activity (2). Recently, a series of phosphonate inhibitors of human ENO, which catalyzes the conversion of 2-PG to PEP in glycolysis and the reverse reaction in gluconeogenesis, have been developed as potential treatments for cancer in the brain (5). These compounds have been shown to cure a rodent model of GBM through inhibition of mammalian ENO2 while being well-tolerated in non-human primate models.

### Phosphonohydroxamates inhibit *N. fowleri* ENO *in vitro*

*N. fowleri* harbors a single ENO gene (*NfENO,* NF0118810) that is 47% identical to human ENO2 (Table 1), being predicted to be a 512 amino acid protein (molecular weight, 56 kDa). (Please note, the ENO gene from the Ty strain of *N. fowleri*, *NfTy_054390*, encodes a protein predicted to be 662 residues, due to an extension at the N-termini but otherwise identical). In both genes, the N-terminal region is predicted to be disordered; a truncation of NF0118810 (residues 44-512) yielded soluble protein, while full-length protein expression was not successful. This truncation, NafoA.00379.a.B5.PW38978, had an apparent *K*_M_ for 2-PG of 923 ± 0.336 µM, within the ranges reported for human skeletal muscle ENO (0.2 mM) and *Leuconostoc mesenteroides* (2.61 mM) (26, 27); a similar truncation used in crystallography studies yielded PDB 7UGH.

**Table 1.** Comparison of the sequences of human and amoebae ENO proteins.

|  | *Nf*ENO | *Bm*ENO |
| --- | --- | --- |
| *Nf*ENO | 100 | 47/63 |
| *Bm*ENO | 47/63 | 100 |
| ENO2 | 47/63 | 73/84 |

Numbers are % identity and % positives
*Bm*ENO (BamaA005554t1; TRINITY\_DN5271\_c0\_g1\_i3\_-\_ORF\_2\_(frame\_2))
ENO2 (*ENOG\_Human*, P09104)

*NfENO* shares key residues required for interactions with phosphonate inhibitors found to inhibit human ENO2, including Arg372 (in ENO2) required to form a salt bridge with the compound (5). Additionally, the limited cytotoxicity and promising distribution to the brain suggested that the phosphonate compounds could be ideal early lead chemotypes for anti-*N. fowleri* development. As a first step to assess the possible use of the agents against amoebae, we tested a panel of analogs against *Nf*ENO and the human enolases (Table 2). The phosphonate compounds were potent inhibitors of recombinant *Nf*ENO. SF2312, a phosphonate antibiotic produced by the actinomycete *Micromonospora* with pan-enolase inhibition activity (6), had an IC_50_ value of 0.31 ± 0.07 µM. HEX (1-hydroxy-2-oxopiperidin-3-yl) phosphonic acid, a six-membered ring structure, had improved specificity for ENO2 (5) and similarly was the most potent of the compounds tested against *Nf*ENO with an IC_50_ value of 0.14 ± 0.04 µM. As it is against ENO2, HEX is a competitive inhibitor of *Nf*ENO. Five- and seven-membered analogs of HEX were active against *Nf*ENO, including deoxy-SF2312 and HEPTA (Table 2). Modification of the phosphonates to generate prodrugs largely eliminated enzyme inhibition, likely due to steric interference of the added pivaloyloxymethyl (POM) groups. BenzylHEX, which was designed as a control that cannot bind (and therefore cannot inhibit) human ENO2 (5), also had no impact on *Nf*ENO at 10 µM, indicating the hydroxamate portion of HEX is important for activity.

**Table 2.** Phosphonate ENO2 inhibitors inhibit *Nf*ENO and have anti-amebic activity.

| Compound | Structure | *Nf*ENO<br>IC<sub>50</sub> (μM) | *Hs*ENO2<br>IC<sub>50</sub> (μM) | *N. fowleri*<br>EC<sub>50</sub> (μM) |
| --- | --- | --- | --- | --- |
| SF2312 |  | 0.31 ± 0.07 | 0.63 ± 0.01 | 1.6 ± 0.11 |
| POMSF |  | > 10 | >10 | 0.66 ± 0.09 |
| HEX |  | 0.14 ± 0.04 | 0.36 ± 0.04 | 0.21 ± 0.02 |
| BenzylHEX |  | > 10 | >10 | > 10 |
| POMHEX |  | > 10 | >10 | 0.70 ± 0.10 |
| CDP25 |  | > 10 | ND | 0.31 ± 0.15 |
| Deoxy-SF2312 |  | 1.2 ± 0.47 | 0.10 ± 0.04 | 0.95 ± 0.20 |
| Benzyl- deoxy-SF2312 |  | >10 | >10 | >10 |
| HEPTA |  | 4.7 ± 1.4 | 5.8 ± 0.88 | >10 |
| Benzyl-HEPTA |  | >10 | >10 | >10 |

ND – Not determined; Bn – -CH<sub>2</sub>-C<sub>6</sub>H<sub>5</sub>

To assess the selectivity of the compounds, inhibitors were tested against human ENO2 (gamma-enolase, *ENO2*). Generally, the sensitivities of ENO2 to inhibitors paralleled those of *Nf*ENO, with HEX being the most potent ENO2 inhibitor tested (Table 2). The one notable difference was deoxy-SF2312, which was more than 10-fold more potent against ENO2 than *Nf*ENO.

### Crystal structure of *Nf*ENO

We determined the X-ray crystal structure of homodimeric *Nf*ENO at 1.95 Å resolution, with 2-PG bound at the active site (PDB 7UGH) (Fig. 1A). A comparison with the human HEX-bound ENO2 (PBD 51DZ (5)) shows that the overall fold is well conserved, with a root-mean-square deviation (RMSD) of 1.98 Å across the 417 Cα atom pairs (Fig. 1B). The phosphate group of the ligands occupies the same position, and the binding pocket is highly conserved; on the superimposed structures, all residues within 5 Å of HEX are identical except for Lys243 that aligns with Ser157 in human ENOs 1, 2, and 3 (see alignment, Fig. S3). In the *Nf*ENO structure, the amine group of the Lys243 side-chain lies close to the phosphate group of the ligand, in contrast to Ser157 in the human enzyme (see Fig. 1B, superposition of the active sites). According to the b-factor, Lys243 is in a flexible loop. Altogether, these observations suggest that Lys243 may be a good target for developing inhibitors specific to the *N. fowleri* enzyme.

**Fig. 1.**
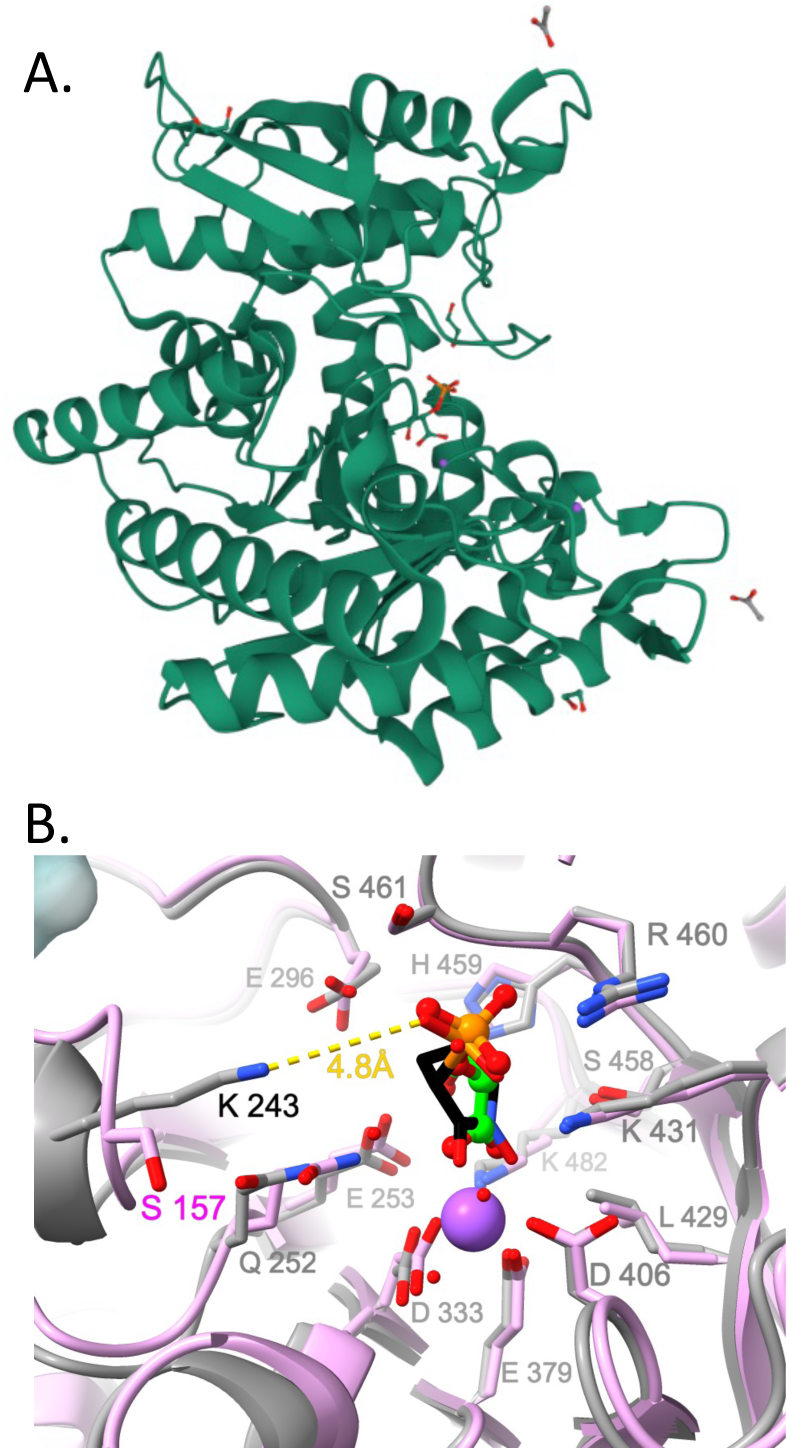
(A) Crystal structure showing 2-PG bound at the active site of *Nf*ENO (PDB 7UGH, 1.95 Å). (B) The active site of *Nf*ENO (PDB 7UGH, grey) overlayed on the human HEX-bound ENO structure (PDB 5IDZ, pink), showing that all residues of the binding pocket are conserved with the sole exception of Lys243, which corresponds to Ser157 in the human enzyme (all other residue numberings refer to *Nf*ENO). Carbon atoms of the *Nf*ENO 2-phosphoglyceric acid ligand are colored green, while those of HEX are shown in black. The phosphate groups of the two ligands overlap and are both positioned close to *Nf*ENO Lys243; the yellow dotted line shows the closest interatomic distance between the Lys243 amine group and the HEX phosphate.

### Molecular modeling of HEX compounds with *Nf*ENO

To gain insight into how HEX, SF2312, deoxy-SF2312, and HEPTA interact with the *Nf*ENO enzyme at the atomic level, we performed a molecular docking simulation of the compounds with *Nf*ENO (PDB 7UGH) using Autodock Vina (Fig. 2). The docking analysis revealed that HEX binds strongly to the active site of *Nf*ENO with a binding affinity of –8.9 kcal/mol, a slightly higher affinity than 2-PG (–8.2 kcal/mol). The binding pose of the *Nf*ENO-HEX complex showed that the hydroxamate and carbonyl moieties chelated a Na^+^ cation while the phosphonate moiety formed polar interactions with Arg460, His459, Ser461, and Ala121 residues. Additionally, Lys243, a residue found in NfENO but not human ENO2, appears to engage in productive interactions with both the hydroxamate and phosphonate through side chain interactions. The *Nf*ENO-HEX interactions from the docking simulation were similar to the interactions observed in the co-crystallization of HEX with human ENO2 (PDB 5IDZ) with the exception of the additional interactions with Lys243 noted above (5). Further, SF2312, deoxy-SF2312, and HEPTA also bind to the active site of the *Nf*ENO enzyme with comparatively weaker binding affinities of –7.6, –6.5, and –6.1 kcal/mol, respectively. Interestingly, Lys243 remains unengaged with bound compound in the docking models of these structural analogs, in contrast to the predicted binding interaction with HEX. These results further underscore the potential importance of leveraging additional interactions with this residue – unique to *Nf*ENO – as a design strategy in future drug development campaigns. The predicted poorer binding affinity of the HEX analogs is consistent with the blunted efficacy of these compounds against *Nf*ENO in vitro as compared to HEX. These results also indicate that the six-membered ring of HEX offers a rather tuned binding conformation to fit into the binding pocket of *Nf*ENO compared with the smaller-sized SF2312 and deoxy-SF2312 and the larger size of HEPTA.

**Fig. 2.**
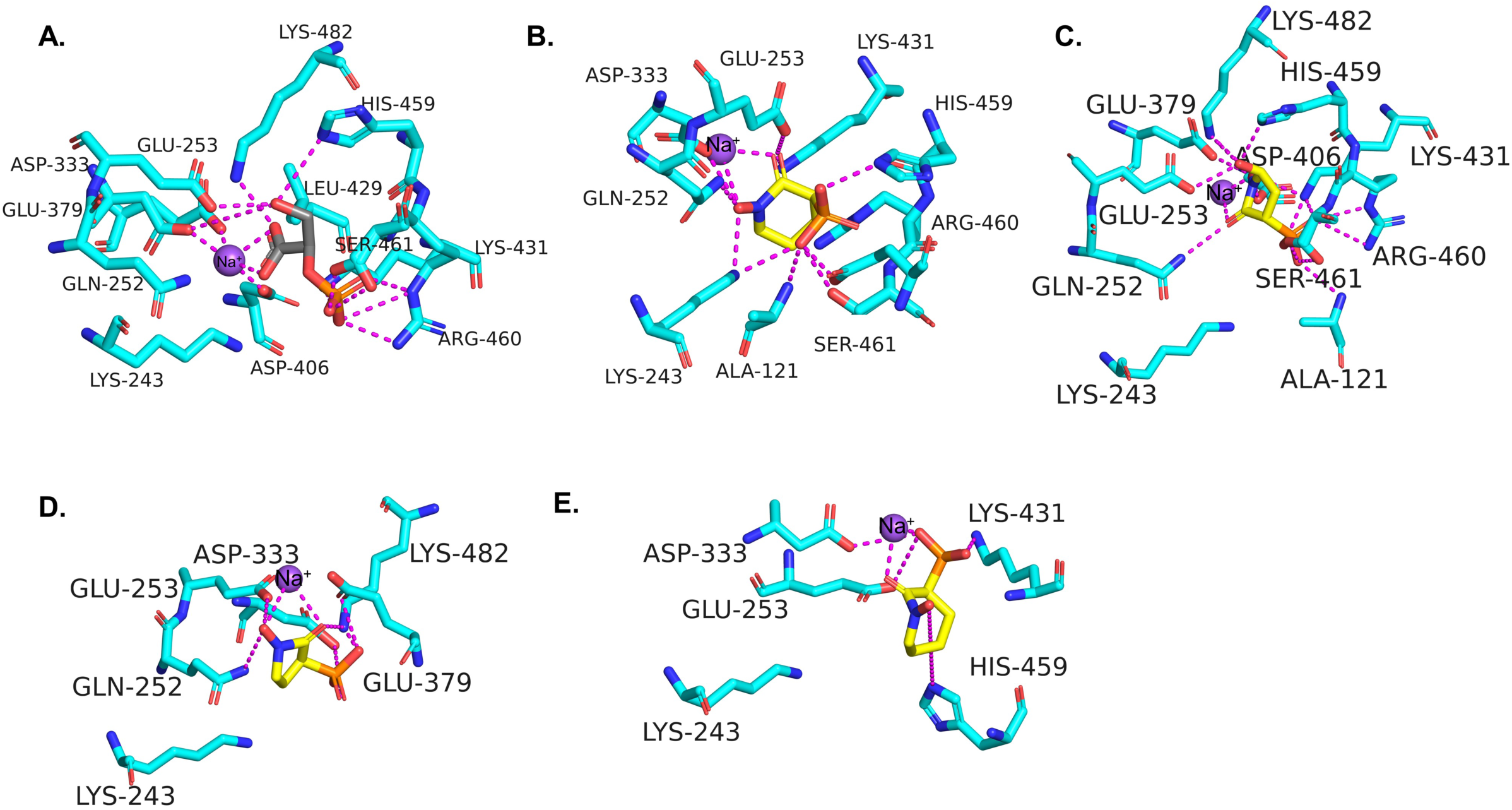
Molecular docking results. Binding poses of (A) 2-PG, (B) HEX (note the unique engagement of Lys243 with drug), (C) SF2312, (D) deoxy-SF2312, and (E) HEPTA at the active site of *Nf*ENO.

### The phosphonate inhibitors are potent anti-amebic compounds

To assess the impact of these compounds on amoebae viability, trophozoites were cultured in the presence of compound or vehicle (1% DMSO) for 48 hr, and viability was scored with CellTiter-Glo reagent. SF2312 was a potent amebicide, with an EC_50_ value of 1.6 ± 0.11 µM (Table 2). Modification of SF2312 to an esterase-labile prodrug by the addition of pivaloyloxymethyl (POM) moieties to yield POMSF modestly (about 2.4-fold) increased the potency of the compound.

HEX was almost 7-fold more potent as an amebicide than SF2312, with an EC_50_ value of 0.21 ± 0.02 μM. Unlike SF2312, modification of HEX with POM moieties to yield POMHEX did not improve the inhibitory potency of HEX. Other phosphonate prodrug analogs, including CDP25, were active in the cell toxicity assays even though they lacked activity against enzyme *in vitro*, suggesting that cellular esterases had processed the compounds into their active form. Nevertheless, none of the prodrugs were as potent as HEX, which was at least six-fold more active against *N. fowleri* than it was against human *ENO1*-deficient cells that it was designed to kill (5). Even a short exposure to HEX was sufficient to suppress amoebae viability, with a one-hour treatment of HEX followed by 48 hr of culture in HEX-free media resulting in an EC_50_ value of 8.06 ± 2.6 µM.

Phosphonate toxicity was not limited to Nf69 strain *N. fowleri* (ATCC 30215) trophozoites, as other strains of *N. fowleri* were also sensitive to the compound, including TY strain *N. fowleri* (ATCC, 30107; EC_50_ value of 0.87 ± 0.23 µM) and Lee strain *N. fowleri* (ATCC 30894) trophozoites maintained on rat neuroblastoma B103 cells (Fig. S4). However, the agent was not broadly anti-amebic, as HEX had no impact on the growth of the non-pathogenic amoebae *N. gruberi*, the pathogenic free-living amoeba *Balamuthia mandrillaris*, or the obligate parasite *Entamoeba histolytica*.

Interestingly, amoebae could be adapted to the presence of HEX by continuous culture in gradually increasing amounts of the agent. Cells grown in the presence of increasing concentration of HEX over a 2.5-month period became more tolerant of the agent, with near-normal growth rates at concentrations ten-fold higher than the EC_50_ value for naïve amoebae. In viability assays, these amoebae were more resistant to the agent, with an EC_50_ value of 0.57 ± 0.30 µM, a value 2.7-fold higher than unacclimated cells in the same medium (0.21 ± 0.03 µM). Removal of selection for three weeks resulted in reversion to HEX sensitivity (EC_50_ value of 0.11 ± 0.01), suggesting the change was dynamic and not permanent.

### Amoebae grown on alternative carbons sources remain sensitive to HEX

Based on transcriptomics analyses, amoebae from the brains of infected rodents upregulate a suite of metabolic genes, including several involved in lipid metabolism (3). We anticipate that *Nf*ENO inhibition would impact both glycolysis (from glucose to pyruvate) and gluconeogenesis (from pyruvate to G6P) (Fig. 3A). However, glycerol from lipid metabolism could circumvent *Nf*ENO inhibition and feed gluconeogenesis through conversion of glycerol 3-phosphate (Gly3P) to dihydroxyacetone phosphate (DHAP) to generate G6P, potentially ameliorating the toxicity of *Nf*ENO inhibition (Fig. 3A).

**Fig. 3.**
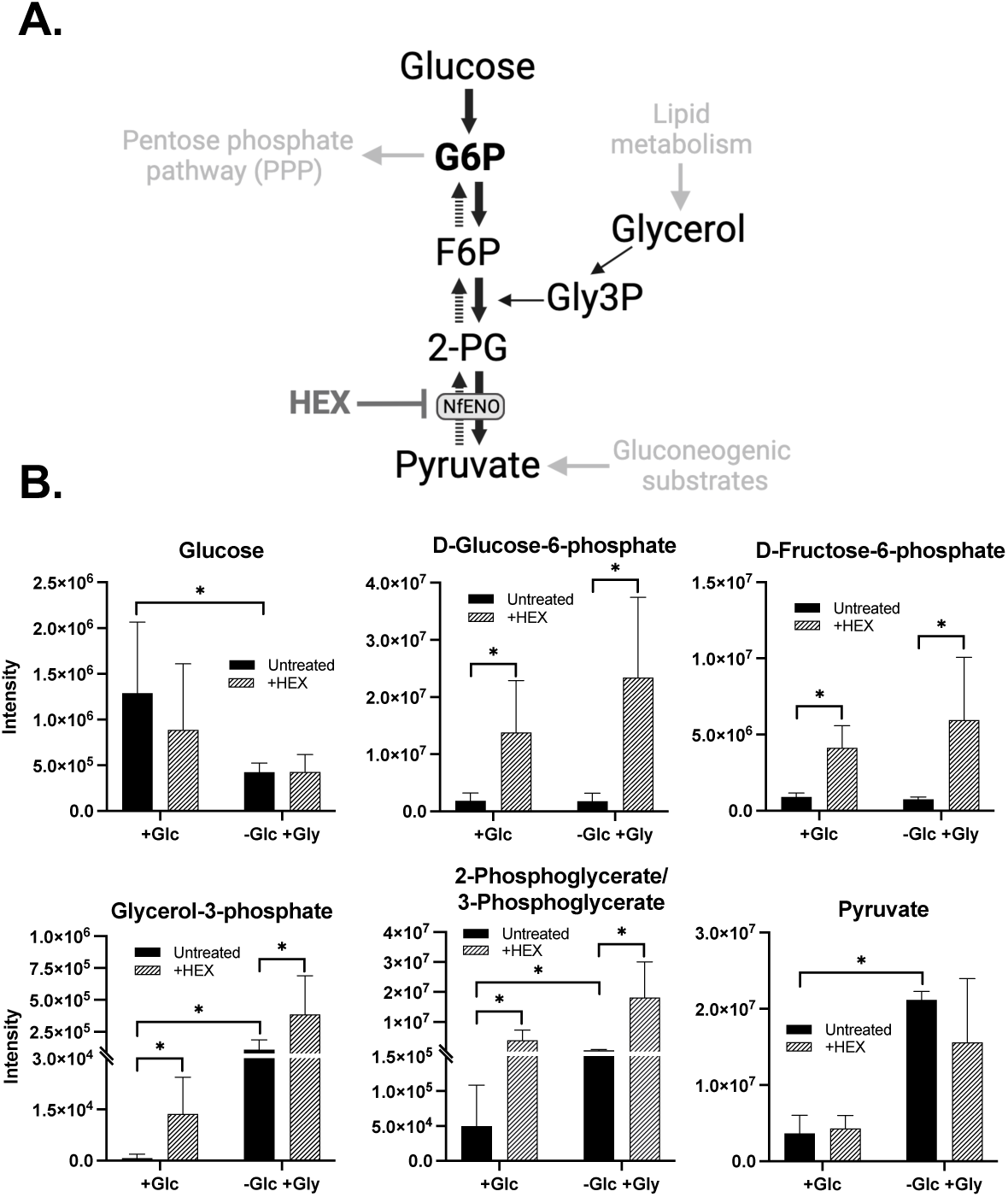
Unbiased metabolomic analysis of glycolytic intermediates after treatment with HEX. A. Schematic of glycolysis and gluconeogenesis in *N. fowleri.* B. Comparative metabolite intensities from amoebae cultured with two different carbon sources (glucose, +glc, or glycerol (-glc,+gly)). Statistical analysis was performed using pairwise two-tail tests with the Benjamini Hochberg method for multiple testing correction with the false discovery rate set at 25%. * indicates significance after the t test and the Benjamini Hochberg correction.

To assess this, amoebae grown in standard medium, which contains abundant glucose (9 mM, “+glc”) were transferred to medium with glycerol (10 mM) and no glucose (“-glc/+gly”), which supported normal trophozoite proliferation with no change in doubling time. While amoebae in glucose-bearing media were highly sensitive to the inhibitor, (EC_50_ values of 0.21 ± 0.02 µM), amoebae cultured in the absence of glucose (with glycerol as an alternative carbon source) were less sensitive to the agent, with an EC_50_ value about 10-fold higher (2.5 ± 0.11 µM).

### HEX treatment *in vitro* leads to accumulation of glycolytic intermediates

Metabolomic profiling of amoebae grown in glucose and treated with HEX revealed a marked accumulation of glycolytic intermediates upstream of *Nf*ENO (Fig. 3B and Supplemental Table S4). The increased intermediates include G6P, F6P, and 2-/3-PG, with 7.4-, 4.5- and 78-fold increases, respectively (see Supplemental Table S5). Pyruvate levels were essentially unchanged. Gly3P abundance, presumably from increased levels of DHAP, also increased 17.9-fold from what was observed in untreated cells.

Amoebae grown in glycerol instead of glucose had an increased abundance of metabolites downstream of the carbon source when compared to cells grown in glucose. Elevated intermediates included Gly3P (150-fold, when compared to cells in glucose), and 2-/3-PG (11-fold). Increased flux from these intermediates may explain the 5.8-fold increase in pyruvate over levels seen in cells grown in glucose. While HEX treatment of these cells led to further accumulation of 2-/3-PG (7.2-fold higher levels than untreated cells grown in glycerol), HEX caused a nearly 50% decrease in pyruvate levels (see Supplemental Table S6).

### HEX extends the life of infected rodents

The promising *in vitro* anti-amebic activity of HEX and its performance in non-human primates, where IP delivery at 300 mg/kg (body weight) was well tolerated and yielded plasma concentrations of about 750 µM for several hours (5), encouraged us to consider the value of the agent in a rat disease model. Rats have a plasma elimination half-life that is very similar to that of the non-human primates (about 60 minutes, (5)), suggesting that parenteral delivery could prove useful. However, disease progression can be rapid, leading us to consider using intranasal instillation as a means of delivery of HEX to the site of infection, including the olfactory bulb.

To assess the impact of HEX on amoebae in an authentic setting, rats were infected by introduction of 2.5 × 10^5^ trophozoites into the nares (Day 0, Fig. 4) and then treatment was initiated two days later by nasal instillation of either HEX or PBS vehicle once a day for 10 days. At the conclusion of treatment on day 11, most (11 of 12) HEX-treated rodents remained. This contrasted with the vehicle treated group, which had five of 12 remaining. By the end of the experiment on the eighteenth day post-infection, 8 of twelve HEX-treated animals remained alive, resulting in an inability to score median survival time, while 1 of twelve vehicle-treated animals survived (median survival time of 10.9 days). Sex did not influence survival after HEX treatment, as there was no difference in median survival time between HEX-treated males and females.

**Fig. 4.**
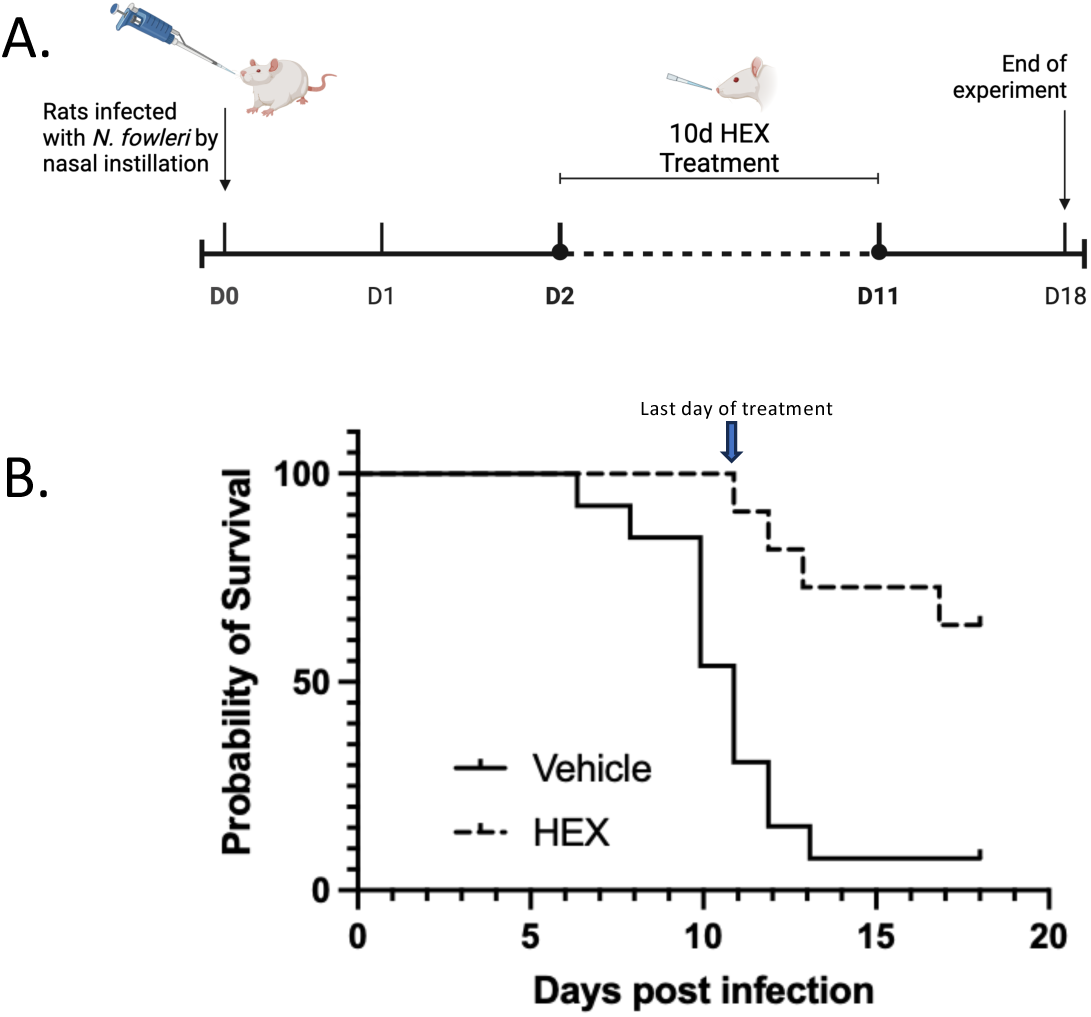
HEX delivered by nasal instillation increased rat median survival of *N. fowleri* infection when compared to vehicle. A. Schematic of experimental set-up. Rats were infected with 2.5 × 10^5^ *N. fowleri* trophozoites on D0 by nasal instillation and then treated with vehicle or HEX for 10 days starting on D2 by nasal instillation. Twelve rats were in each group (6 males, 6 females). B. Kaplan-Meier survival curves for PBS- and HEX-treated groups. Median survival time for vehicle-treated animals was 10.88 days post-infection, while the median survival time for HEX-treated rats was undefined (since >50% of the population survived). A log-rank (Mantel-Cox) test determined a significant difference between the median survival time for each group with a p-value of 0.0006 (***).

Survival could have resulted from either elimination of the pathogen or suppression of the infection. To assess this, brains were extracted from all survivors and scored for infection by culture of the pathogen. Of the eight HEX treated animals that survived the experiment, six had on-going infections as detected by culture of amoebae from extracted brains. The lone PBS treated control survivor was negative for amoebae, suggesting that infection was never established.

## DISCUSSION

Current treatment regimens for *N. fowleri* infection include agents that have been empirically identified because of their use in one of the few cases of successful treatment. More recently, miltefosine and azithromycin have been added to treatment cocktails, but infection outcomes remain poor. With the goal being the development of improved treatments for *N. fowleri* infections, we have explored the value of the amoeba ENO enzyme, a key component in both glycolysis and gluconeogenesis, as a potential therapeutic target. In glycolysis, ENO catalyzes the conversion of 2-PG to PEP for use by pyruvate kinase in the payoff stage that yields ATP. ENO also plays a role in gluconeogenesis, with PEP generated from gluconeogenic substrates like lactate and amino acids being converted by ENO to 2-PG which ultimately yields G6P that is required for the pentose phosphate pathway.

Here, we have found that phosphonate ENO inhibitors designed to be selective for human ENO2 are active against amoeba enzyme and toxic to the trophozoite stage. *N. fowleri* is remarkably sensitive to one such inhibitor, HEX, with EC_50_ values about 6-fold lower than those required to kill ENO-1-deficient GBM cells, the cancer type that was the target of the initial phosphonate design campaign (5). The reasons for the hypersensitivity of the amoebae are unclear but could be related to differences in the uptake of the compound between the protist and human cells. Generally, the cell permeability of phosphate-bearing compounds, including phosphonate-based drugs, is poor given that they are charged at physiological pH (28). Some bacteria use an ABC transporter system encoded in the gene cluster *phnCDE* for phosphonate uptake (29). The *phnC* gene, which shares similarity with nucleotide-dependent permeases, has been identified in eukaryotic phytoplankton that can utilize phosphonates as a phosphorus source (30), and *N. fowleri* harbors a possible homolog (E-value = 2 × 10^-30^, KAF0981242.1, FDP41_013030, AmoebaDB). The amoebae have homologs of additional known phosphonate transporters, although these proteins have other described or predicted roles indicating that phosphonate uptake may be a moonlighting function. These additional candidates include potential glycerol-3-phosphate (Gly3P) and hexose-6-phosphate transporters that have been implicated in the uptake of the phosphonate drugs fosmidomycin and fosfomycin in some bacteria (31), candidates that will be assessed in future loss of function studies as genetic tools for gene manipulation in *N. fowleri* become available.

*N. fowleri* responds to different environments with a dynamic metabolic approach to exploit available resources. *In vitro*, the modest lipid amounts in the 10% fetal bovine serum included in typical media (yielding 2-3% of the fatty acids and cholesterol normally available to cells in a vertebrate (32)) is insufficient to support growth (2). Nevertheless, lipid metabolism may be important to amoebae in the brain, based on observed increases in fatty acid metabolism gene expression in isolates from infected rodents (3). Additionally, lipids are the preferred carbon source of *N. gruberi*. The likely importance of lipid metabolism raises the question of why HEX is toxic to amoebae? First, glucose metabolism remains critical, even in the presence of lipids with glycolytic genes being upregulated in the brain-isolated amoebae. In *N. gruberi*, small levels of glucose are consumed even in the presence of abundant lipids, likely for the critical role of replenishing TCA intermediates through anaplerosis (4).

We anticipate that *Nf*ENO inhibition would impact both glycolysis (from glucose to pyruvate) and gluconeogenesis (from pyruvate to G6P). Given that, glycerol from lipid metabolism could circumvent *Nf*ENO inhibition and feed gluconeogenesis to generate G6P, ameliorating the toxicity of *Nf*ENO inhibition. However, cells grown in either glucose or glycerol remained sensitive to the agent. In both cases, the accumulation of glycolytic intermediates, in particular 2-/3-PG, could explain toxicity. Glycolytic intermediates serve as important allosteric regulators of other pathways, with their accumulation being dysregulating. If this were true, HEX treatment of cells grown in glycerol (which had increased levels of Gly3P, 2-/3-PG, and pyruvate relative to HEX-treated cells in glucose), should have been more sensitive to the agent. But they are not, being about 10-fold less sensitive, suggesting that the causes of HEX toxicity are more complex. One possibility is that 2-PG accumulation disrupts amoebae osmoregulation through an impact on trehalose, an osmoprotectant found in other systems, including *N. gruberi* (33). Trehalose-6-phosphate synthase (TPS, which *N. fowleri* harbors as NfTy_062120) is required for trehalose synthesis and could be inhibited by 2-PG (34). While we anticipate that HEX-triggered accumulation of 2-PG would inhibit TPS similarly in cells grown in either glycerol or glucose, the glycerol in the medium may partially substitute for trehalose as an osmoprotectant.

While HEX did not eliminate the amoebae in most of the treated animals, it did have a profound impact on the development of disease, with only one rodent succumbing to infection during treatment. This suggests that higher concentrations of agent or increased frequency of treatment could prove useful for eradication of the infection. Of note, HEX resistance was not a likely explanation for the lone HEX treatment failure, as amoebae isolated from the HEX treated rat remained as sensitive to HEX as the amoebae from a vehicle treated rodent (Fig. S5).

While most of the treated animals remained infected, disease symptoms were largely suppressed while HEX was being provided. It is unclear how the amoebae were avoiding the lethal agent, but it is possible that in the brain environment the organisms encyst, lying quiescent until treatment was removed. Nevertheless, the delay in disease progression may prove useful in treatment regimens, as HEX may allow other promising, but slow acting, anti-amebic agents time to exert their lethal effect. Further, continued treatment beyond the 10-day period could sufficiently suppress the amoebae to allow the host immune system to respond to eliminate the invader.

Intranasal instillation has proven to be a fast-acting route of drug administration for several brain-active agents, including anticonvulsive compounds (35, 36). The success of the HEX treatment may be in part due to direct transport of the agent to the brain, which would reduce the impact of dilution by distribution (37). Direct transport can result from endocytosis by olfactory sensory cells followed by exocytosis into the synaptic clefts in the olfactory bulb, which would introduce HEX into the CNS in a position to interact with invading amoebae at the site of brain infection (38).

It is unlikely that HEX will serve as a component of a pan-species anti-amebic cocktail as the compound is inactive against other pathogenic free-living amoebae, including *Balamuthia mandrillaris. B. mandrillaris* harbors an ENO (*Bm*ENO, 000586F| quiver) that is 56% identical to human ENO2 and is predicted to have the residues required for HEX binding (Table 1). The lack of sensitivity of *B. mandrillaris* suggests that *Bm*ENO may be less important for the metabolic success of the amoebae. Alternatively (or additionally), the observed difference in toxicity may result from differences between the amoebae in HEX transport or metabolism.

In this study, we have found that inhibition of central carbon metabolism can impact amoebae viability, both *in vitro* and in an animal model. The favorable pharmacological profiles of HEX and related phosphonates suggests that they may be useful in the treatment of *N. fowleri* brain infections (39), and their promising activity in a rodent model supports this possibility. Ultimately, the ENO inhibitors may prove useful for future therapeutics, either alone or in concert with other agents (like lipidolysis inhibitors) that would in tandem block multiple key pathways required for both ATP and reducing equivalent biosynthesis with catastrophic consequences.

## Supporting information

Supplemental materials, including Tables and Figures

## ACKNOWLEDGMENTS

The authors would like to thank Drs. Dennis Kyle and Christopher Rice for their support and Drs. Heather Walters and Lesly Temesvari for their assistance in testing HEX against *Entamoeba.* We thank Dr. Paul Leonard (MD Anderson Cancer Center Core for Biomolecular Structure and Function) for providing purified recombinant human enolases. The authors would also like to thank Colm Roster and Caroline Palmentiero for their technical support in these projects. Work from JCM’s laboratory is partially supported by R21 AI175463. Additionally, the JCM and DCW laboratories were supported in part by the NIH Center for Biomedical Excellence (COBRE) grant under award numbers P20GM146584 and P20GM113226. SMW was supported by R01 CA231509. The SSGCID has been funded by Federal funds from the National Institute of Allergy and Infectious Diseases, National Institutes of Health, Department of Health and Human Services, under Contract Nos.: 75N93022C00036, HHSN272201200025C, HHSN272200700057C and HHSN272201700059C.

## The abbreviations used are

2-PG, 2-phosphoglycerate; 2-/3-PG, a mixture of 2-PG and 3-phosphoglycerate; DHAP, dihydroxyacetone phosphate; ENO2, human enolase 2; F6P, fructose 6-phosphate; G6P, glucose 6-phosphate; G6PDH, glucose 6-phosphate dehydrogenase; GBM, glioblastoma multiforme; Gly3P, glycerol 3-phosphate; HEPTA, (1-hydroxy-2-oxoazepan-3-yl) phosphonic acid; HEX, (1-hydroxy-2-oxopiperidin-3-yl) phosphonic acid; *Nf*ENO, *N. fowleri* enolase; SF2312, (1,5-dihydroxy-2-oxopyrrolidin-3-yl) phosphonic acid

