## Supplemental materials, including Tables and Figures for "Enolase inhibitors as therapeutic leads for *Naegleria fowleri* infection"

**List of Supplementary Materials**

**Materials and Methods**

**Protein purification, crystallization, and structure determination**

An *E.coli* codon optimized construct in the vector pQE-30 was purchased from Twist Bioscience containing residues 2-512 of the full length 512 amino acid protein (NF0118810, AmoebaDB, https://amoebadb.org/amoeba/app/). Sequence comparison and secondary structure predictions using Xtalpred (https://xtalpred.godziklab.org/XtalPred-cgi/xtal.pl) were used to design multiple constructs to test for optimal crystallization construct. Five constructs composed of residues 14-512, 28-512, 44-512, 62-512, 80-512 were processed using the SSGCID high throughput pipeline (*20-22*). The constructs were cloned using into the ligation independent cloning (LIC) (*37*) expression vector pBG1861 (*21*) encoding a non-cleavable 6xHis fusion N-terminal tag. Protein was expressed in *E. coli* BL21(DE3)R3 Rosetta cells and two constructs yielded soluble protein, constructs bearing residues *Nf*ENO(44-512) and *Nf*ENO(62-512). These lines were used for large scale (2L) protein production in auto-induction media (*23*) in a LEX Bioreactor (Epiphyte Three Inc.), as previously described (*22*). The expression clones NafoA.00379.a.B4.GE44253 (NfENO(62-512) and NafoA.00379.a.B5.GE44254(NfENO(44-512) are available at https://www.ssgcid.org/available-materials/ssgcid-proteins/.

*Nf*ENO(44-512) and *Nf*ENO(62-512) were purified in a two-step protocol consisting of $Ni^{2+}$-affinity and size-exclusion chromatography (SEC). All chromatography was performed on an ÄKTApurifier 10 (GE Healthcare) according to previously described procedures (*20*). Thawed bacterial pellets were lysed by sonication in lysis buffer (25 mM HEPES pH 7.0, 500 mM NaCl, 5% glycerol , 0.5% CHAPS, 30 mM Imidazole, 10 mM $MgCl_2$, 1 mM TCEP, 250 ug/ml AEBSF, and 0.025% sodium azide). After sonication, the crude lysate was clarified by mixing with 500 units of Benzonase (RT, 45 min). The lysate was clarified by centrifugation and then passed over a Ni-NTA His-Trap FF 5 ml column (GE Healthcare) which was pre-equilibrated with loading buffer (25 mM HEPES pH 7.0, 500 mM NaCl, 5% glycerol , 30 mM imidazole, 1 mM TCEP, and 0.025% sodium azide). The column was washed and protein pooled and concentrated prior to loading on a SEC column (Superdex 75, GE Healthcare) equilibrated with running buffer (loading buffer without imidazole). The SEC peak fractions eluted as a single large peak in the molecular-mass range ~50 kDa, suggesting monomeric enzyme which was consistent with the size of the denatured purified protein, as determined by SDS-PAGE. Peak fractions were pooled and concentrated using an Amicon purification system (Millipore) to a final concentration of to 73 mg/ml *Nf*ENO(44-512) and 45 mg/ml *Nf*ENO(62-512).

Purified *Nf*ENO(44-512) and *Nf*ENO(62-512) were screened for crystallization in 96-well sitting-drop plates against the JCSG++ HTS (Jena Bioscience) and PACT premier HT96 (Molecular Dimensions) crystal screens. Proteins were diluted in SEC buffer 1:1 to final concentrations 36 mg/ml *Nf*ENO(44-512) and 23 mg/ml *Nf*ENO(62-512). Equal volumes of protein and precipitant solutions were set up at 289 K against reservoir in sitting-drop vapor-diffusion format. In addition to protein-only crystallizations, co-crystallization trials with 2-phosphoglycerate (2-PG, 5 mM) and HEX (5 mM) were also pursued. Only NfENO(44-512) in complex with 2-PG crystallized to form diffraction quality crystals in 85 mM Tris/HCl pH 8.5, 25.5% (w/v) PEG4000, 15% glycerol, and 170 mM sodium acetate, 5 mM 2-PG. The crystals were flash-frozen by plunging directly into liquid nitrogen without cryoprotectant exchange.

X-ray data were collected at 100°K on beamline 21-ID-F, LS-CAT, at the Advanced Photon Source, Argonne National Laboratory. Data were processed with XDS reduced with XSCALE (*24*) and the structure was solved by molecular replacement using the structure of *B. subtilus* ENO (PDB 4A3R) as a search model (*25*). Structures were refined using iterative cycles of Phenix (*26*) followed by manual rebuilding of the structure using Coot (*27*) to yield a resolution of 1.95Å, with the quality of all structures checked using MolProbity (*28*). All data-reduction and refinement statistics are shown in Tables S4 and S5. Structure figures were prepared and analyzed using PyMOL (v.1.5; Schrodinger) and coordinates and structure factors have been deposited with the Protein Data Bank www.rcsb.org (*29*) with accession number 7UGH.

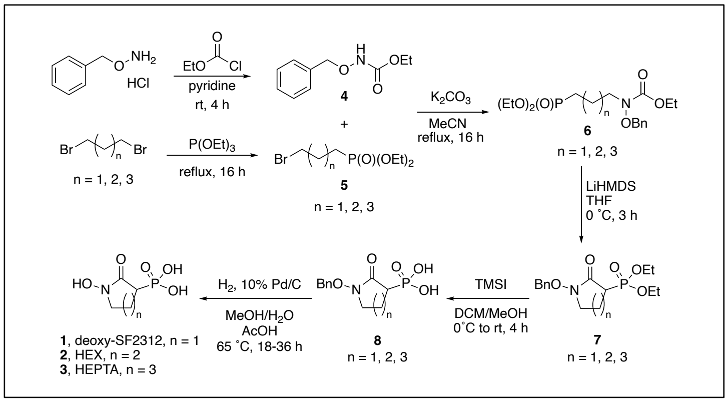

**Fig S1.** Synthetic route for deoxy-SF2312, HEX, and HEPTA. The compounds (1-hydroxy-2-oxopyrrolidin-3-yl) phosphonic acid (deoxy-SF2312) **1**, (1-hydroxy-2-oxopiperidin-3-yl) phosphonic acid (HEX) **2**, and (1-hydroxy-2-oxoazepan-3-yl) phosphonic acid (HEPTA) **3**, were synthesized following published procedures reported in the literature (*5, 8*). Briefly, a mixture of *O*-benzylhydroxylamine hydrochloride and chloroformate in pyridine was stirred at room temperature under $N_2$ for 4 h to yield ethyl benzyloxycarbamate (**4**). Triethyl phosphite in excess of the appropriate dibromoalkane (*i.e.* n = 1 for Deoxy-SF2312, n = 2 for HEX, and n = 3 for HEPTA) was stirred at 90 ˚C overnight to yield the appropriate alkylbromophoshonate (**5**) as light yellow oil after purification by silica gel column chromatography (0–8% MeOH in DCM). Potassium carbonate was added to a solution of **4** and **5** in MeCN and the mixture was stirred at 90 ˚C overnight to yield the linear diethyl phosphonate-carbamate (**6**) as a yellow oil after purification by silica gel column (0–8% MeOH in DCM). Treatment of **6** in THF with LiHMDS at 0 ˚C for 3 h under $N_2$ afforded the cyclized diethyl phosphonate (**7**) as a yellow oil after purification by silica gel column chromatography (0–8% MeOH in DCM). Hydrolysis of **7** with iodotrimethylsilane in DCM at 0–25 ˚C for 4 h under $N_2$ yielded the benzyloxy phosphonic acid (**8**) as a yellow solid after reverse phase column chromatography (C18 silica gel; 0–25% MeOH in $H_2O$). Finally, the benzyl group was removed by hydrogenolysis of (1.8 mmol, **8**) dissolved in 10 mL of $H_2O$/MeOH (1:1 v/v) containing 30 mmol of acetic acid and palladium on carbon (10%, 50 mg) stirred at 65 ˚C for 18–36 h at 1 atm $H_2$ to give the desired products as an off-white oil (**1**) or light-yellow oil (**2** and **3**).

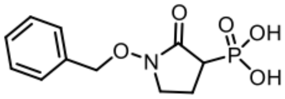

**(1-(benzyloxy)-2-oxopyrrolidin-3-yl)phosphonic acid**; yellow solid; **$^{1}$H-NMR** (500 MHz, DMSO) δ 7.44-7.36 (m, 5H), 4.88 (s, 2H), 3.38 (m, 1H), 3.33 (m, 1H), 2.71 (m, 1H), 2.12 (m, 2H); **$^{13}$C NMR** (125 MHz, DMSO) δ 167.1 (d, J=4.3 Hz, 1C), 135.4, 129.1 (s, 2C), 128.5, 128.3 (s, 2C), 75.5, 44.8 (d, J=4.2 Hz, 1C), 37.9, 17.9 (d, J=3.7 Hz, 1C); **$^{31}$P ($^{1}$H decoupled) NMR** (200 MHz, DMSO) δ 18.6 (s, 1P).

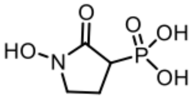

**(1-hydroxy-2-oxopyrrolidin-3-yl)phosphonic acid**; off-white oil; **$^{1}$H-NMR** (500 MHz, $D_2O$) δ 3.65 (m, 1H), 3.57 (m, 1H), 2.93 (m, 1H), 2.31 (m, 1H); 2.22 (m, 1H); **$^{13}$C NMR** (125 MHz, $D_2O$) δ 168.4, 47.4 (d, J=4.3 Hz, 1C), 38.9 (d, J=137.1 Hz, 1C), 17.4 (d, J=3.5 Hz, 1C); **$^{31}$P ($^{1}$H decoupled) NMR** (200 MHz, $D_2O$) δ 19.9 (s, 1P).

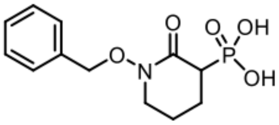

**(1-(benzyloxy)-2-oxopiperidin-3-yl)phosphonic acid**; yellow solid; **$^{1}$H-NMR** (500 MHz, $D_2O$) δ 7.49-7.42 (m, 5H), 4.94 (q, J=6.4, 10.3, 10.5 Hz, 2H), 3.49 (m, 2H), 2.96 (dt, J=25.0, 6.5 Hz, 1H), 2.02 (m, 1H), 1.94 (m, 1H), 1.78 (m, 1H); **$^{13}$C NMR** (125 MHz, $D_2O$) δ 166.2 (d, J=5.3 Hz, 1C), 134.3, 129.9 (s, 2C), 129.1, 128.7 (s, 2C), 75.7, 50.1, 43.0 (d, J=128.8 Hz, 1C), 22.1 (d, J=3.8 Hz, 1C), 21.2 (d, J=7.9 Hz, 1C). **$^{31}$P ($^{1}$H decoupled) NMR** (200 MHz, $D_2O$) δ 19.7 (s, 1P).

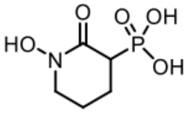

**(1-hydroxy-2-oxopiperidin-3-yl)phosphonic acid**; light yellow oil; **$^{1}$H-NMR** (500 MHz, $D_2O$) δ 3.65 (m, 2H), 3.00 (m, 1H), 2.12 (m, 1H), 1.88 (m, 1H), 1.70 (m, 1H); **$^{13}$C NMR** (125 MHz, $D_2O$) δ 165.9 (d, J=6.2Hz, 1C), 51.5, 40.9 (d, J=130.7 Hz, 1C), 21.8 (d, J=3.6 Hz, 1C), 20.9 (d, J=8.1 Hz, 1C). **$^{31}$P ($^{1}$H decoupled) NMR** (200 MHz, $D_2O$) δ 20.4 (s, 1P).

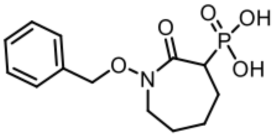

**(1-(benzyloxy)-2-oxoazepan-3-yl)phosphonic acid**; yellow solid; **$^{1}$H-NMR** (500 MHz, DMSO) δ 7.45-7.32 (m, 5H), 4.85 (s, 2H), 3.80 (m, 1H), 3.76 (m, 1H), 2.89 (m, 1H), 2.02-1.89 (m, 2H), 1.62-1.54 (m, 4H); **$^{13}$C NMR** (125 MHz, DMSO) δ 168.6 (d, J=3.3 Hz, 1C), 135.7, 129.2 (s, 2C), 128.4, 128.3 (s, 2C), 75.4, 64.9, 51.2, 44.7 (d, J=138.5 Hz, 1C), 26.4, 15.2. **$^{31}$P ($^{1}$H decoupled) NMR** (200 MHz, D2O) δ 19.0 (s, 1P).

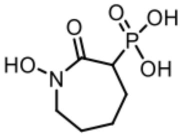

**(1-hydroxy-2-oxoazepan-3-yl)phosphonic acid**; light yellow oil; **$^{1}$H-NMR** (500 MHz, DMSO) δ 3.82 (dd, J = 15.4, 6.9 1H), 3.72 (dd, J=15.8, 8.7 Hz, 1H), 2.89 (ddd, J=22.6, 9.1, 2.5 Hz 1H), 1.95 (m, 1H), 1.85 (m, 1H), 1.58-1.42 (m, 4H); **$^{13}$C NMR** (125 MHz, DMSO) δ 171.9 (d, J=3.1 Hz, 1C), 63.1, 51.2, 41.0 (d, J=138.3 Hz, 1C), 26.5, 14.1. **$^{31}$P ($^{1}$H decoupled) NMR** (200 MHz, DMSO) δ 16.0 (s, 1P).

Solvent DMSO
MHz 500
Nucleus $^{1}H$

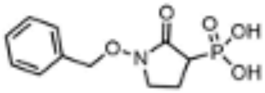

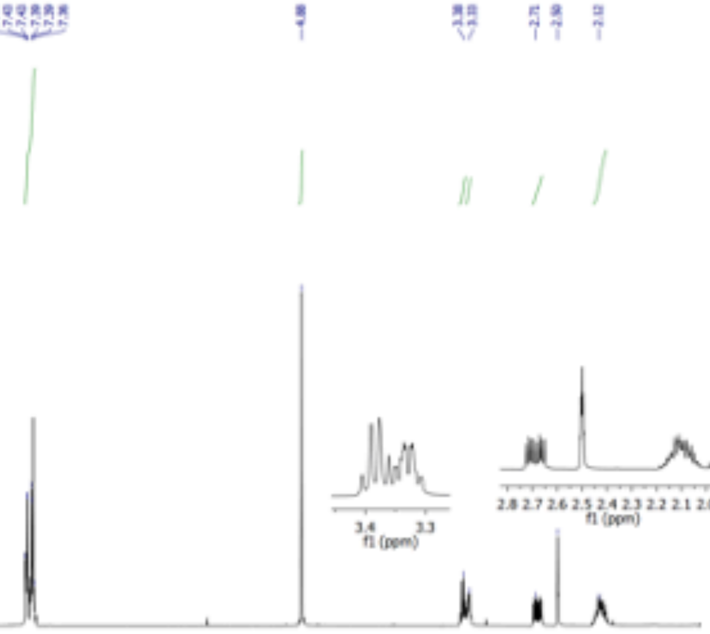

Solvent DMSO
MHz 125
Nucleus $^{13}C$

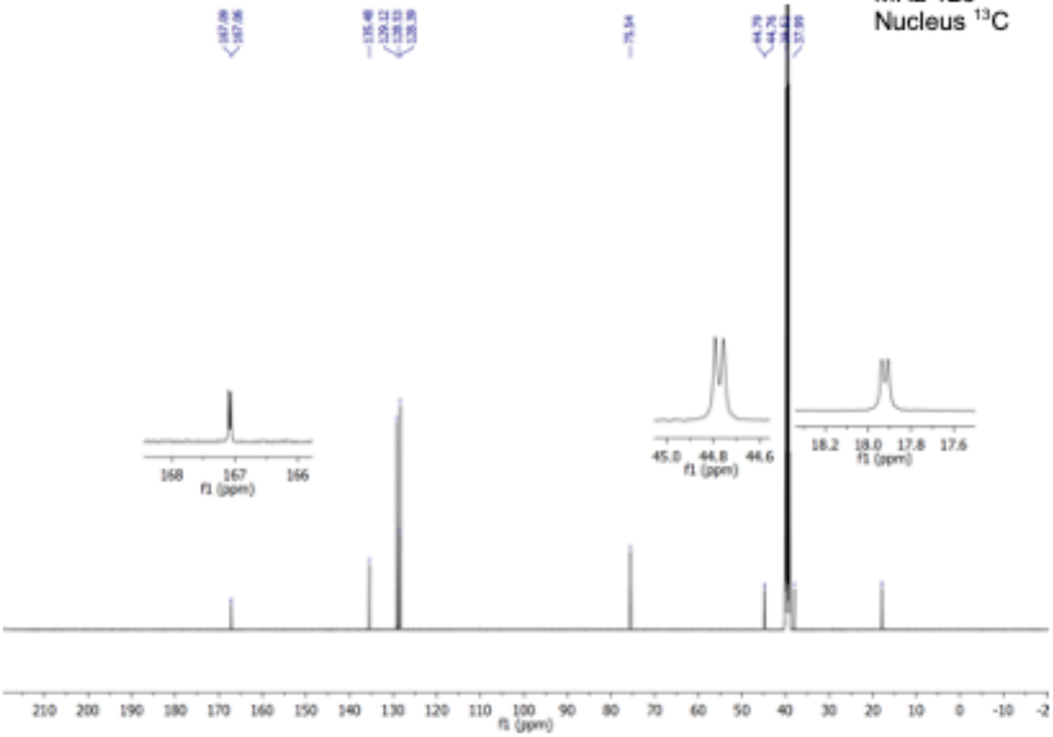

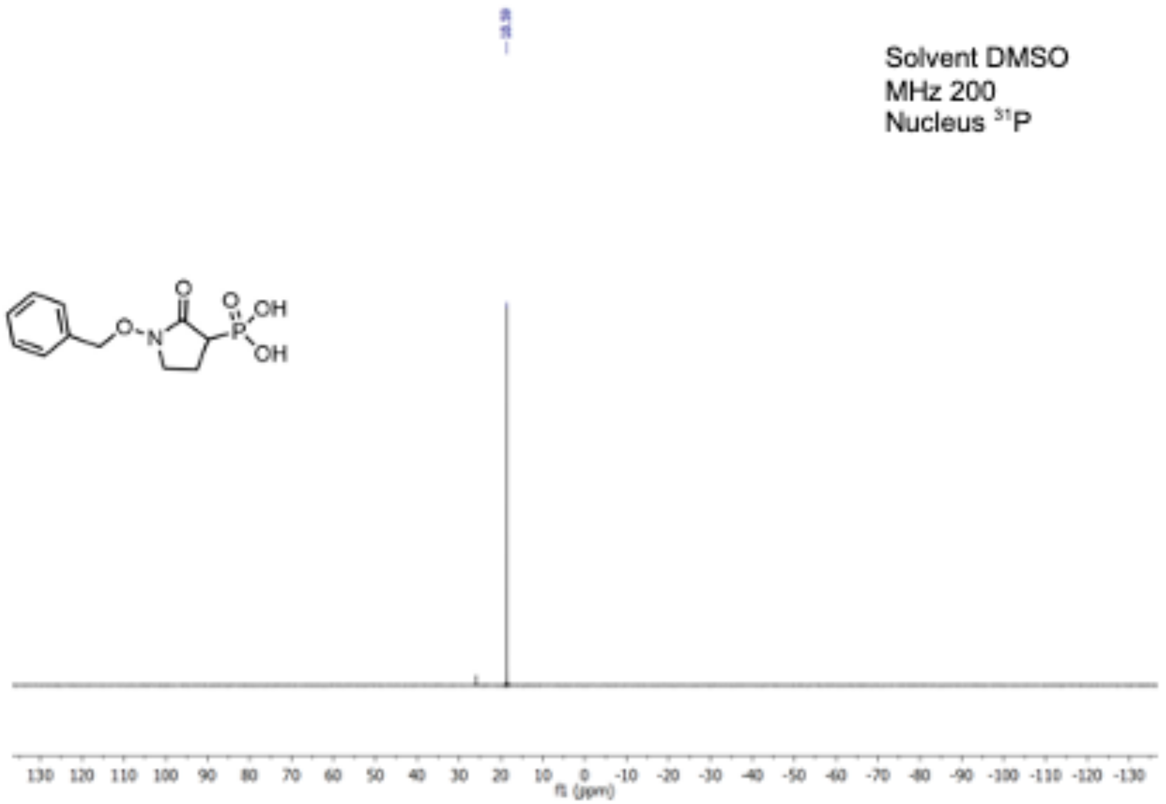
Solvent DMSO
MHz 200
Nucleus 31P
O
O
OH
OH
N
P
130 120 110 100 90 80 70 60 50 40 30 20 10 0 -10 -20 -30 -40 -50 -60 -70 -80 -90 -100 -110 -120 -130
f1 (ppm)

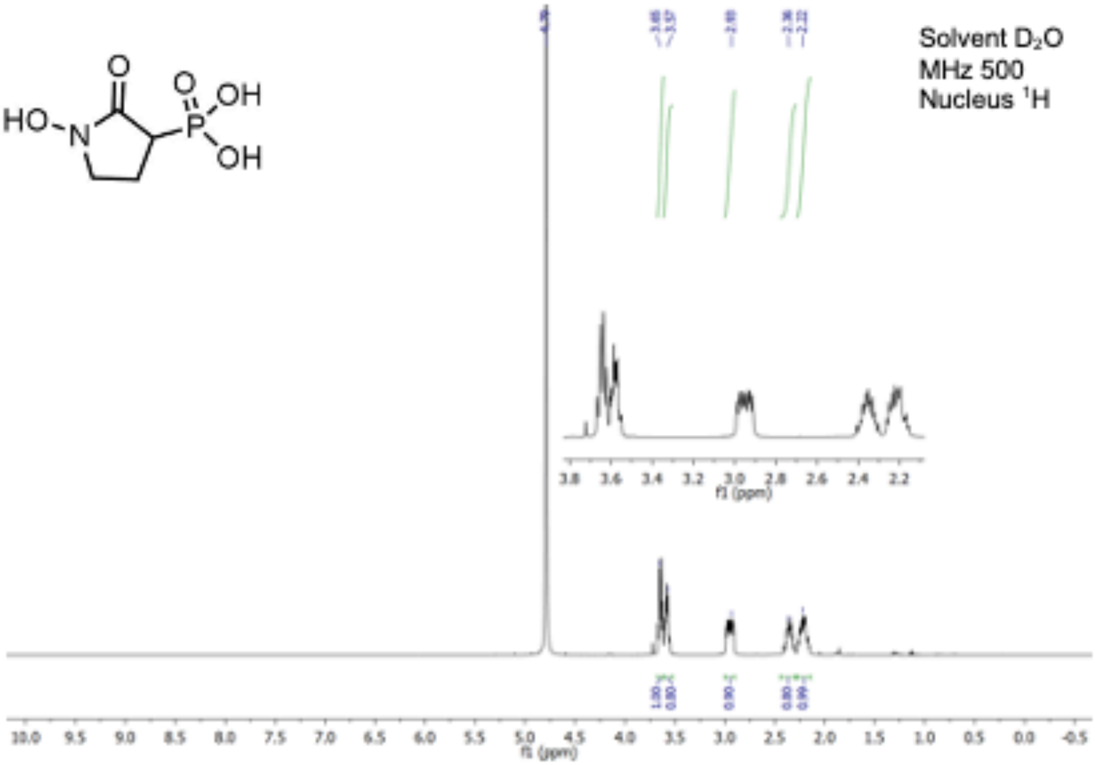

Solvent $D_2O$
MHz 500
Nucleus $^1H$

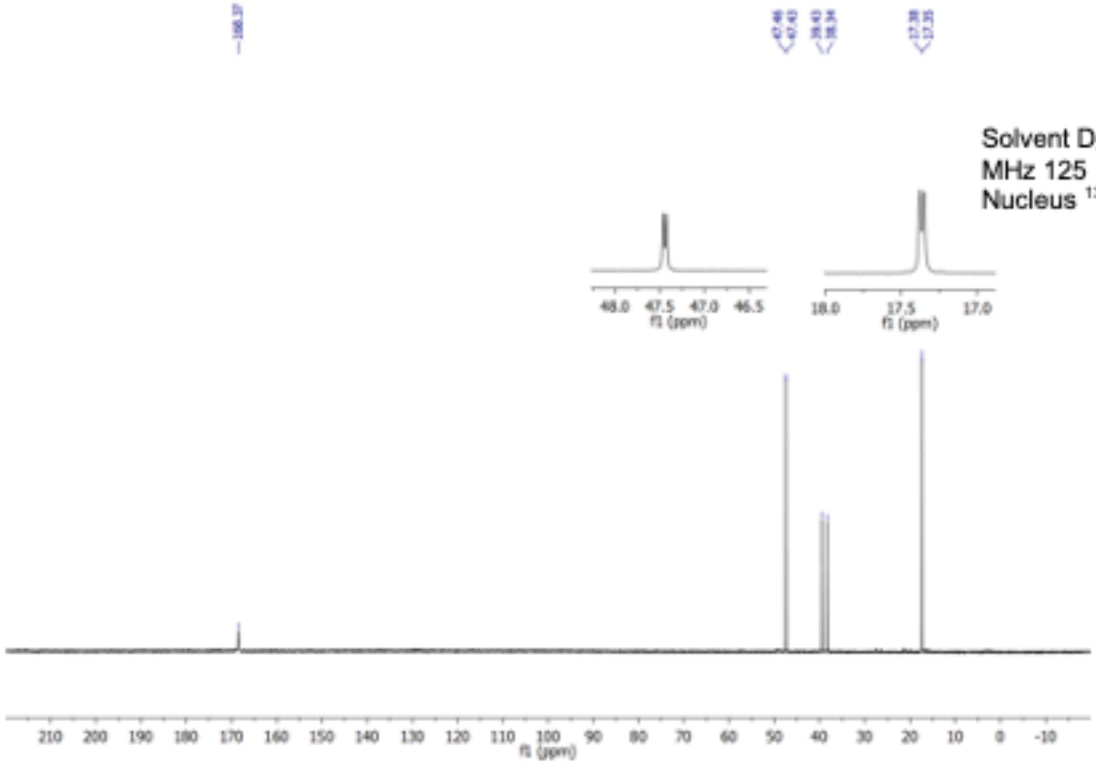

Solvent $D_2O$
MHz 125
Nucleus $^{13}C$

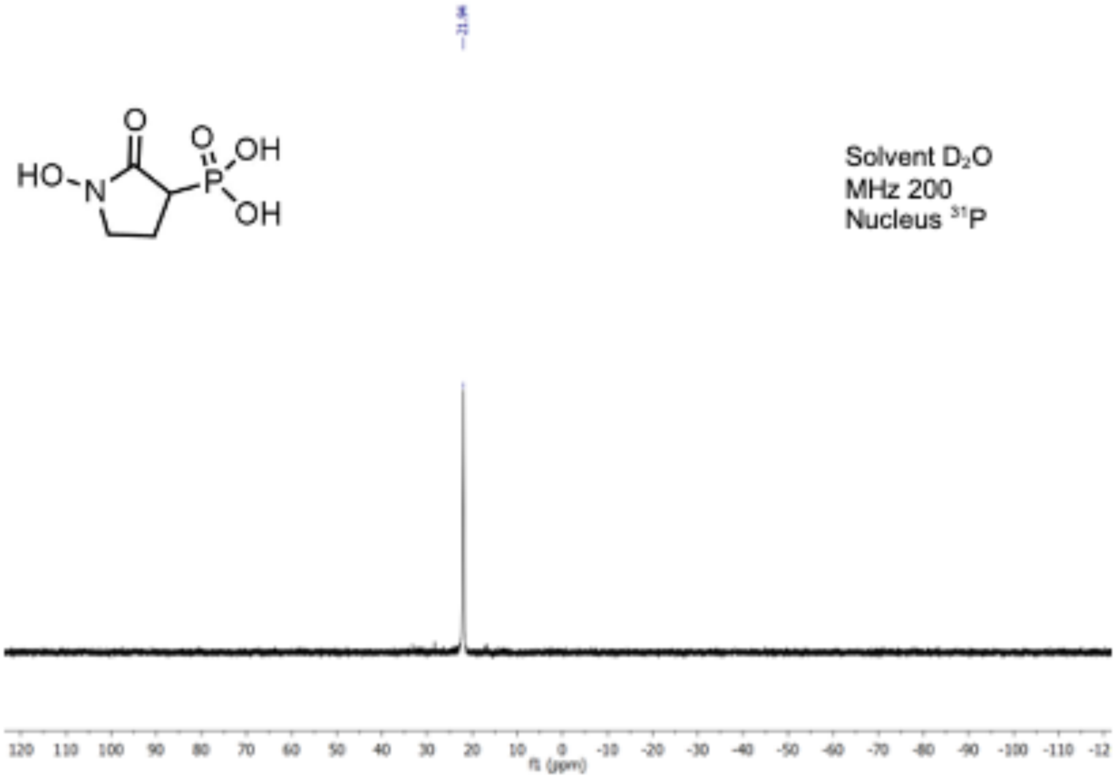
HO
N
O
P
OH
OH
Solvent D2O
MHz 200
Nucleus 31P
120 110 100 90 80 70 60 50 40 30 20 10 0 -10 -20 -30 -40 -50 -60 -70 -80 -90 -100 -110 -12
f1 (ppm)

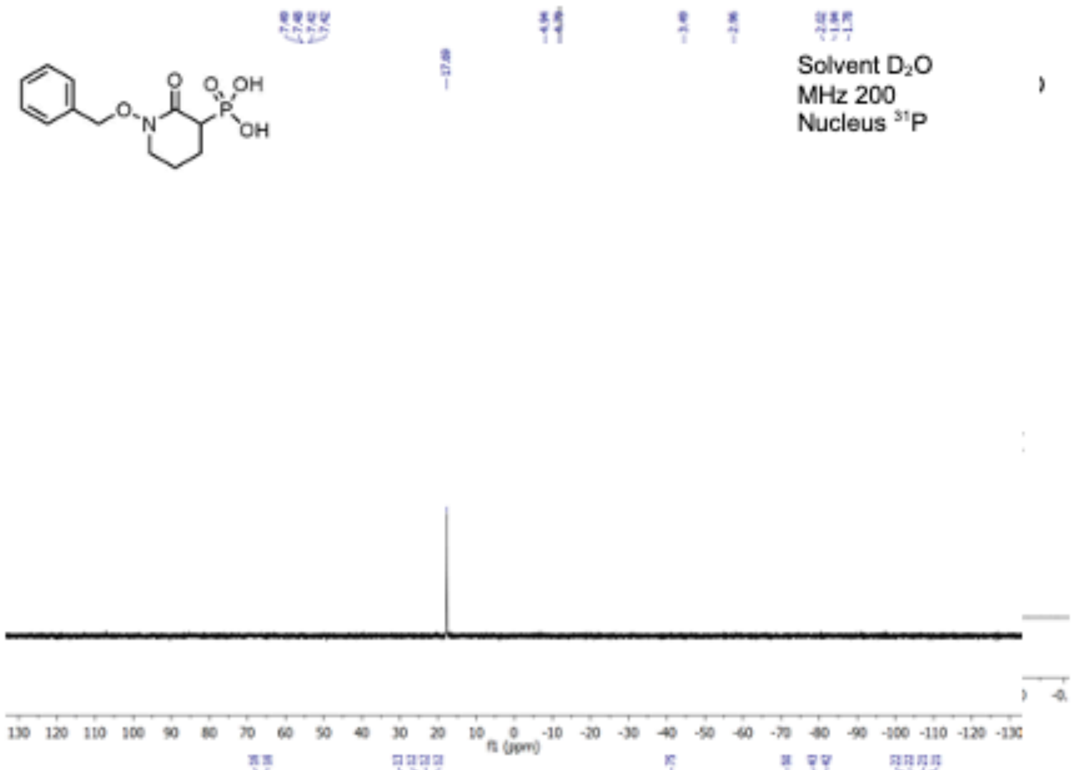

Solvent D2O
MHz 200
Nucleus 31P
f1 (ppm)

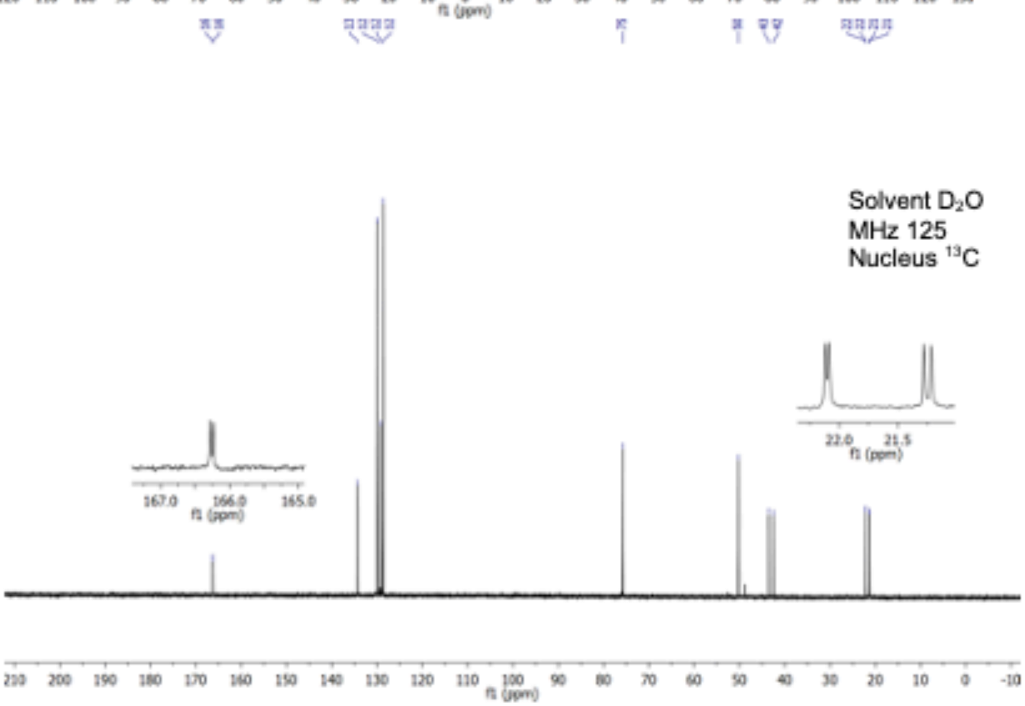

Solvent D2O
MHz 125
Nucleus 13C
f1 (ppm)

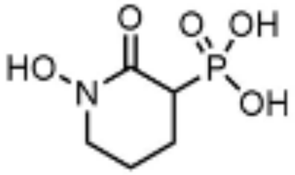

Solvent $D_2O$
MHz 200
Nucleus $^{31}P$

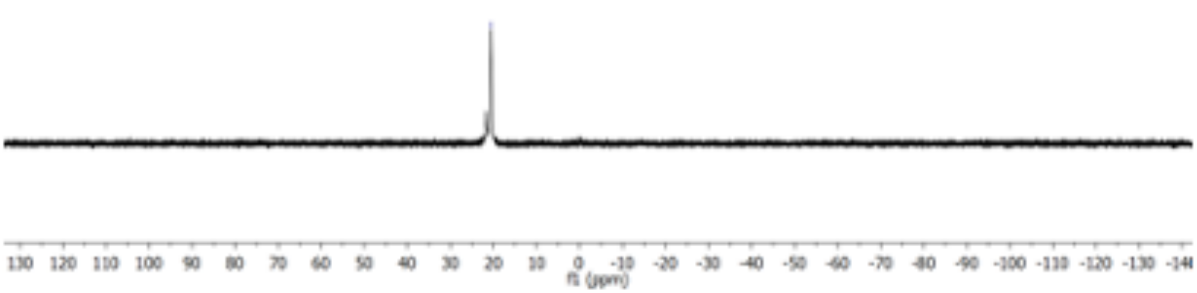

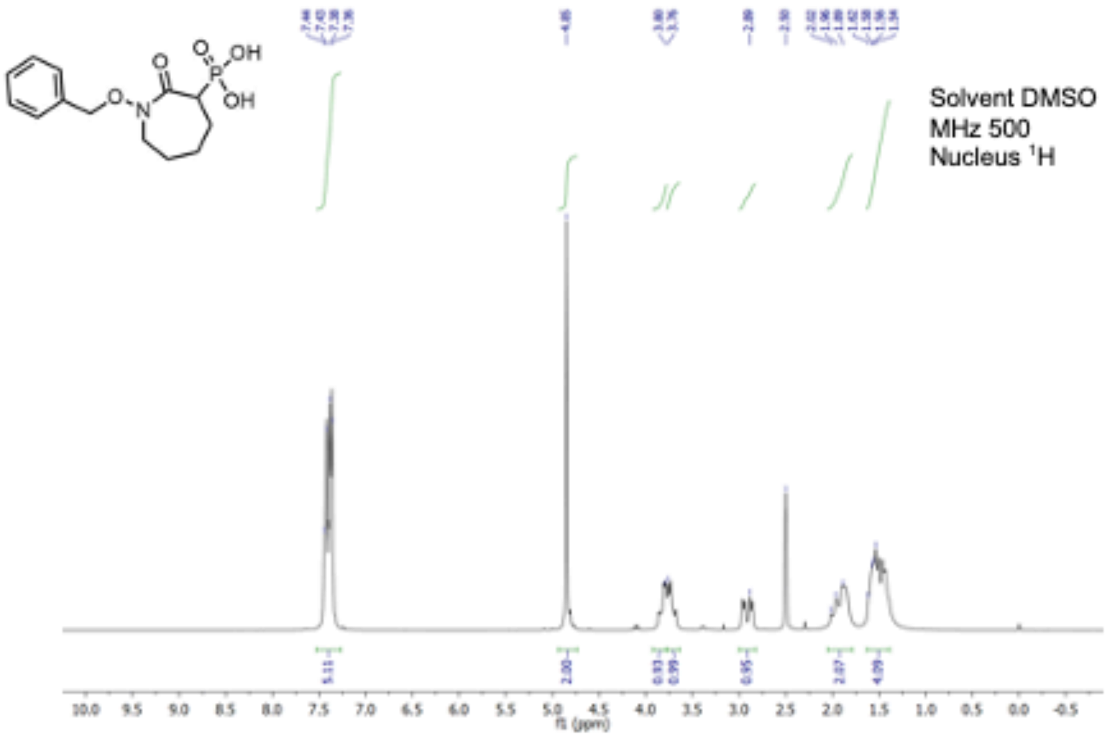
Solvent DMSO
MHz 500
Nucleus ¹H
f1 (ppm)

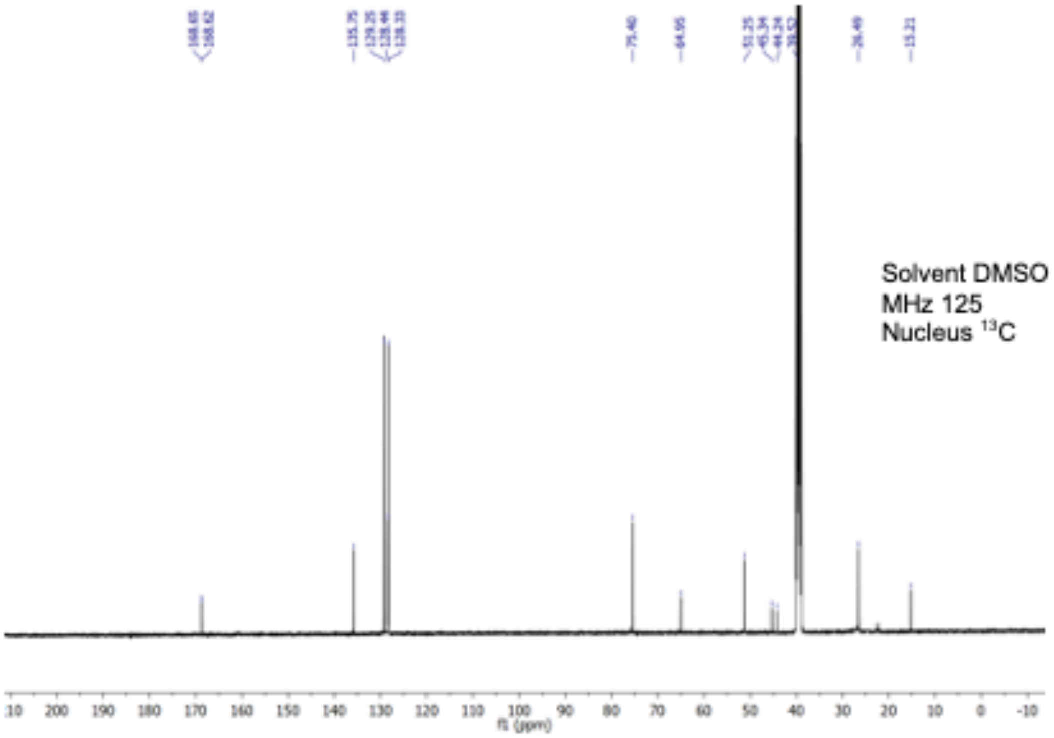
Solvent DMSO
MHz 125
Nucleus ¹³C
f1 (ppm)

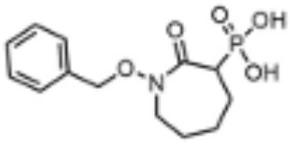

Solvent DMSO
MHz 200
Nucleus $^{31}P$

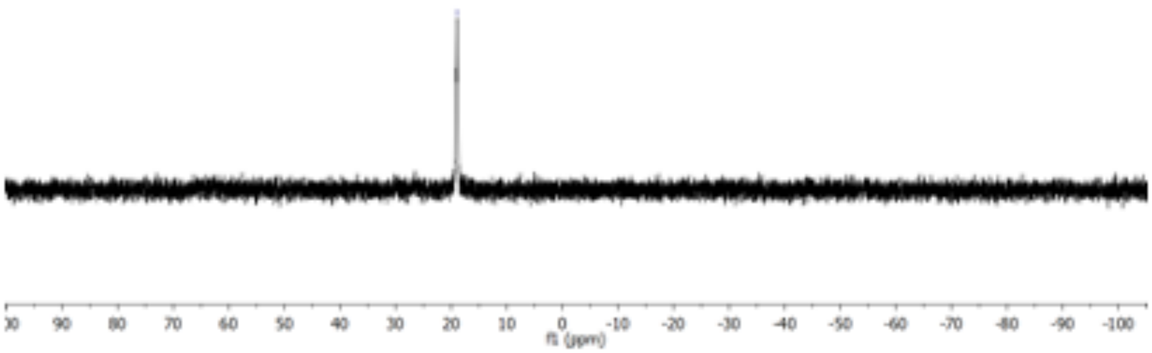

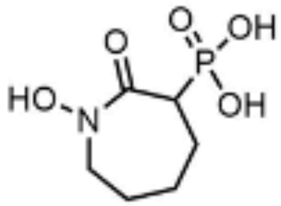

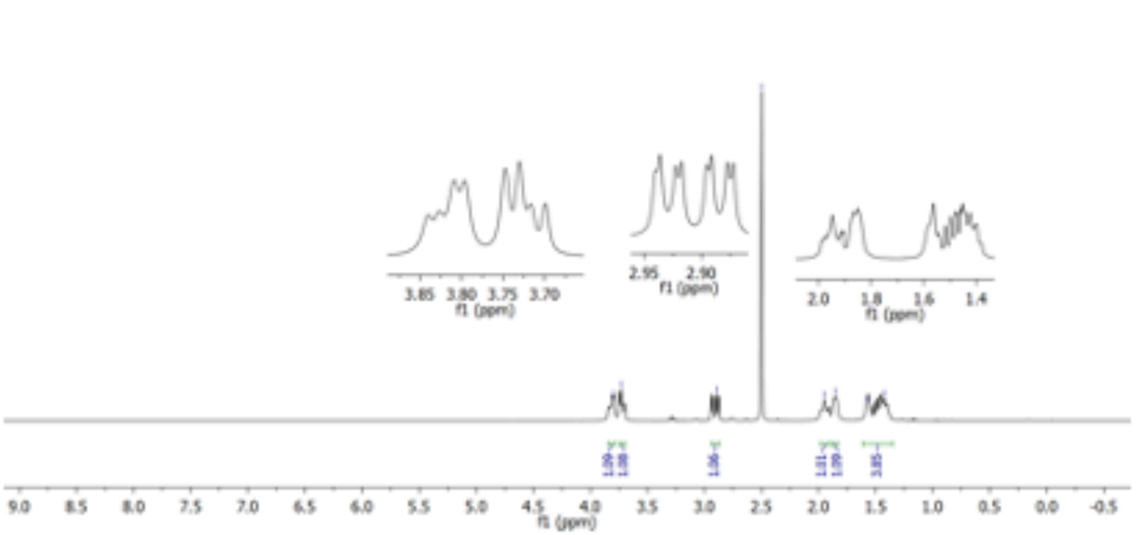
Solvent DMSO
MHz 500
Nucleus ¹H
f1 (ppm)

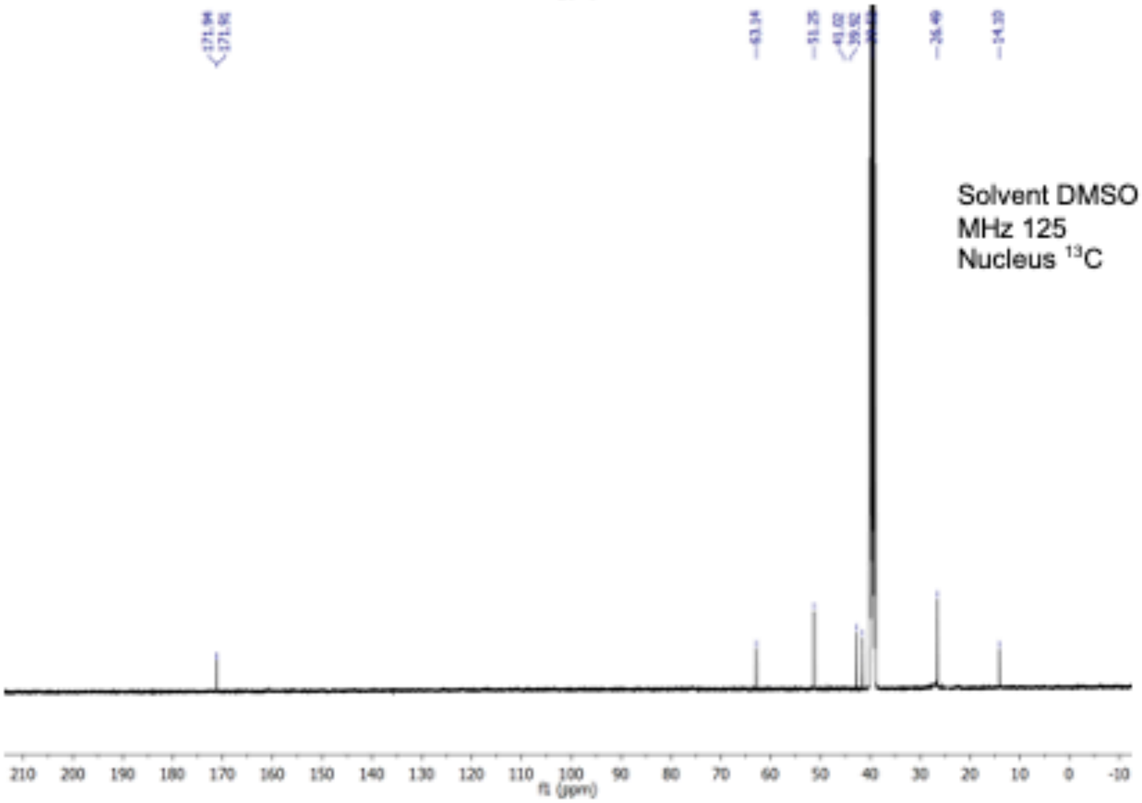
Solvent DMSO
MHz 125
Nucleus ¹³C
f1 (ppm)

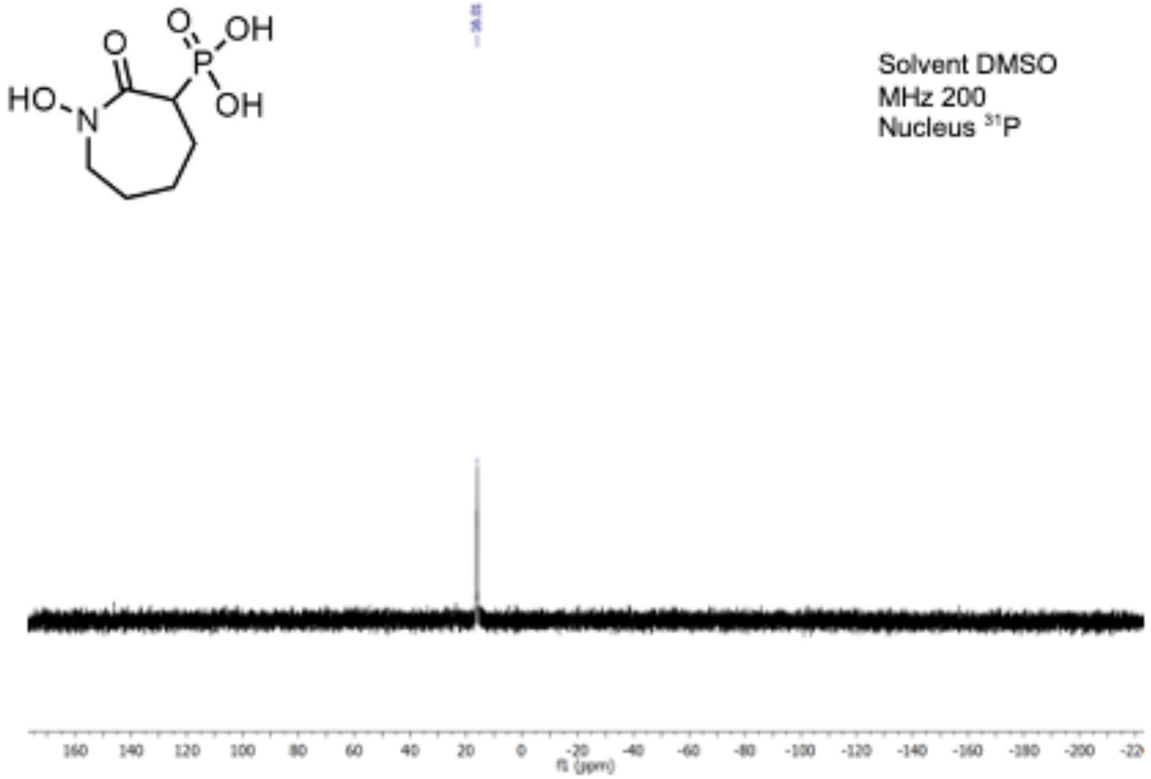

**Fig S2.** Spectroscopic characterization of deoxy-SF2312, HEX and HEPTA - $^{1}H$ and $^{13}C$ NMR spectra were obtained using Bruker Advance 500 MHz spectrometers. Chemical shifts are reported in parts per million (ppm). Spectra are referenced to residual solvent peaks. Flash column chromatography was carried out using ZEOCHEM silica gel (40-63 μm). Analytical and preparative thin-layer chromatography (TLC) were performed on Sorbtech silica G TLC plates. All non-aqueous reactions were performed under an inert atmosphere of nitrogen in flame-dried glassware containing a stir bar unless otherwise noted. Acetonitrile (ACN), tetrahydrofuran (THF), dichloromethane (DCM), methanol (MeOH), and pyridine were obtained from commercial sources and dried following standard distillation procedures. All other solvents were obtained from commercial sources and used without drying unless otherwise noted. All water and aqueous solutions were made using deionized (DI) water.

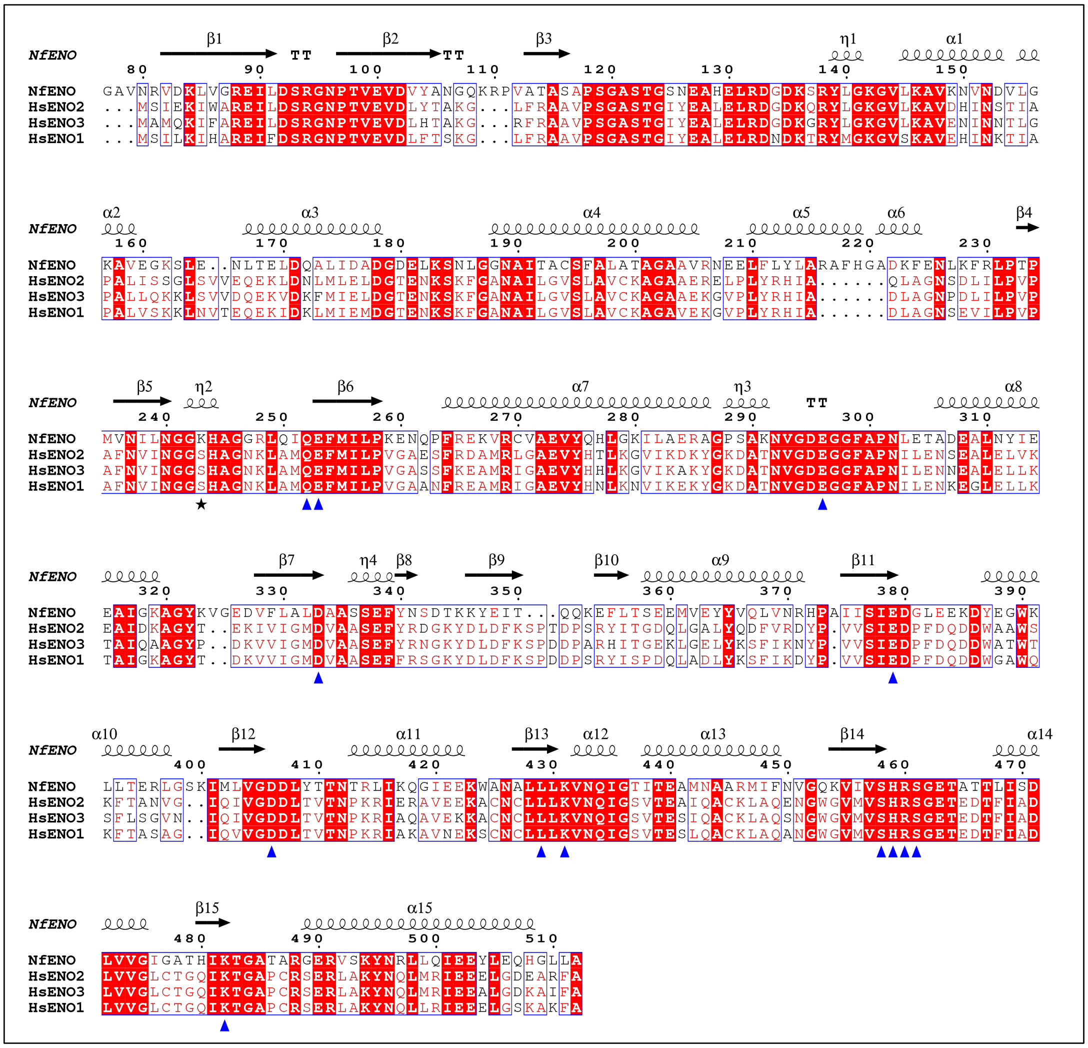

**Fig S3.** ClustalOmega alignment of *Nf*ENO with human ENOs colored by percent equivalent score [https://doi.org/10.1093/nar/gku316]. The top lines show the secondary structure elements of *Nf*ENO. The scale above the alignment corresponds to the *Nf*ENO sequence. Active site residues are highlighted with a blue triangle; the black star marks the position of *Nf*ENO Lys243.

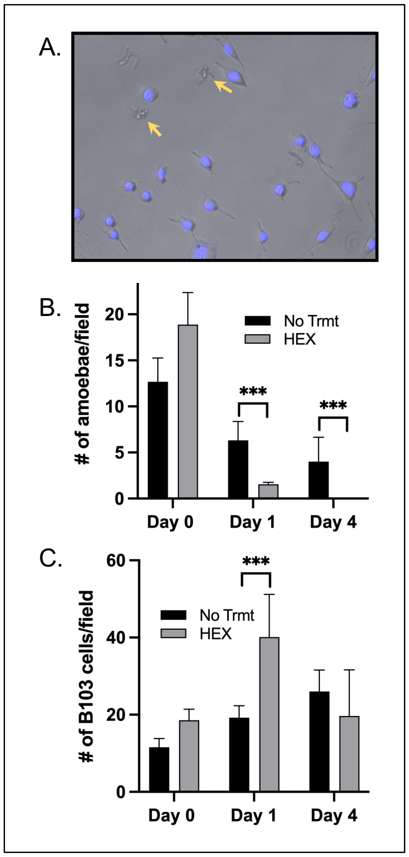

**Fig S4.** Lee strain *N. fowleri* grown on feeder cells were sensitive to HEX. (A.) Lee strain amoebae (1 x $10^4$ cells) were seeded into 96-well plate wells that were near-confluent with rat neuroblastoma B103 cells in DMEM and NCM at a 1:1 ratio and visualized after staining with Hoechst 33342 Solution (Thermo Fisher Scientific) for 10 min. Amoeabe are indicated by yellow arrows. (B.) HEX is toxic to Lee strain trophozoites. Amoebae and feeder cells were plated, and after 24 hours, spent medium was removed and 100 µM HEX was added with fresh medium. After an additonal 0, 24, or 96 hrs, cells were stained with Hoechst 33342 Solution and amoeabe were counted, with at least four fields of cells counted from each of three replicate wells. (C.) Impact of 100 µM HEX on rat neuroblastoma B103 cells. To assess feeder and amoeabe cell viability, Hoechst stained nuclei were counted, or trophozoites were scored by visualization, respectively. At least four fields of cells from each of three replicate wells were counted for both. The amoebae were readily distinguished from the host cells by the large difference in nuclei size. *** indicates $p < 0.001$.

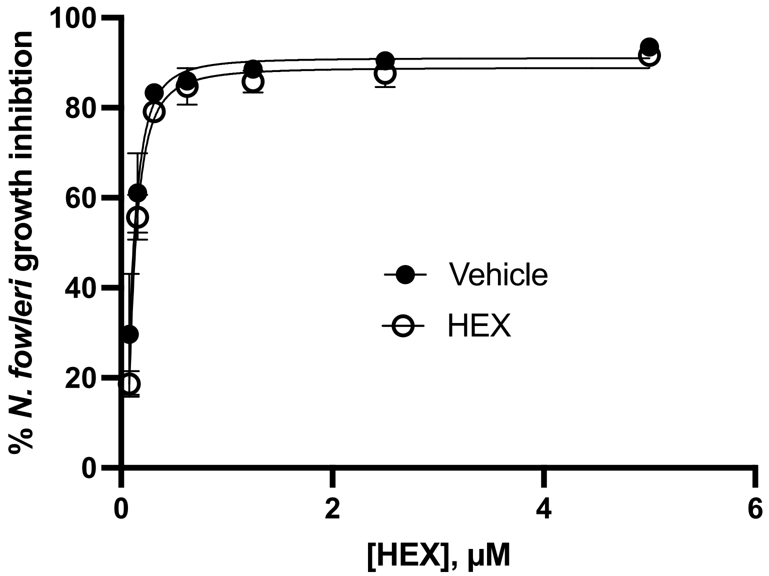

**Fig S5.** Amoeba resistance to HEX does not explain the therapeutic failure observed in the one HEX-treated rodent that succumb to infection. Trophozoites were cultured from brains of a HEX-treated (HEX) or PBS-treated (Vehicle) rodent for two weeks in media followed by testing against HEX in a standard viability assay. Both cultures responded similarly, with $EC_{50}$ values of 0.09 ± 0.04 and 0.1 ± 0.08 µM for HEX- or Vehicle-treated rodents, respectively. Drug concentrations were tested in triplicate, with some error bars smaller than the graphing symbols.

**Supplemental Table S1. NfENO sequences from Uniprot, AmoebaDB, and the truncation used for crystallography that is deposited in the PDB.**

| Source | ID | Sequence |
|---|---|---|
| Uniprot | tr\|A0A6A5BXC3\|<br>A0A6A5BXC3_N<br>AEFO | MTDKQPVSEYLQNHNLQKLVEDALNECYNANASDPVGFLGHFFLNRGKKGAVNRVDKLV<br>GREILDSRGNPTVEVDVYANGQKRPVATASAPSGASTGSNEAHELRDGDKSRYLGKGVL<br>KAVKNVNDVLGKAVEGKSLENLTELDQALIDADGDELKSNLGGNAITACSFALATAGAAVR<br>NEELFLYLARAFHGADKFENLKFRLPTPMVNILNGGKHAGGRLQIQEFMILPKENQPFREK<br>VRCVAEVYQHLGKILAERAGPSAKNVGDEGGFAPNLETADEALNYIEEAIGKAGYKVGED<br>VFLALDAASSEFYNSDTKKYEITQQKEFLTSEEMVEYYVQLVNRHPAIISIEDGLEEKDY<br>EGWKLLTERLGSKIMLVGDDLYTTNTRLIKQGIEEKWANALLLKVNQIGTITEAMNAARM<br>IFNVGQKVIVSHRSGETATTLISDLVVGIGATHIKTGATARGERVSKYNRLLQIEEYLEQ<br>HGLLA |
| AmoebaDB | NF0118810 | MLYLVLMLVENGFGQNCFLHSQKICSNMTDKQPVSEYLQNHNL**Q**KLVEDALNECYNANA<br>SDPVGFLGHFFLNRGKKGAVNRVDKLVGREILDSRGNPTVEVDVYANGQKRPVATASAP<br>SGASTGSNEAHELRDGDKSRYLGKGVLKAVKNVNDVLGKAVEGKSLENLTELDQALIDAD<br>GDELKSNLGGNAITACSFALATAGAAVRNEELFLYLARAFHGADKFENLKFRLPTPMVNIL<br>NGGKHAGGRLQIQEFMILPKENQPFREKVRCVAEVYQHLGKILAERAGPSAKNVGDEGG<br>FAPNLETADEALNYIEEAIGKAGYKVGEDVFLALDAASSEFYNSDTKKYEITQQKEFLTSEE<br>MVEYYVQLVNRHPAIISIEDGLEEKDYEGWKLLTERLGSKIMLVGDDLYTTNTRLIKQGIEE<br>KWANALLLKVNQIGTITEAMNAARMIFNVGQKVIVSHRSGETATTLISDLVVGIGATHIKTGA<br>TARGERVSKYNRLLQIEEYLEQHGLLA |
| PDB (truncation used for crystallography) | 7UGH_1 | MAHHHHHHQKLVEDALNECYNANASDPVGFLGHFFLNRGKKGAVNRVDKLVGREILDSR<br>GNPTVEVDVYANGQKRPVATASAPSGASTGSNEAHELRDGDKSRYLGKGVLKAVKNVN<br>DVLGKAVEGKSLENLTELDQALIDADGDELKSNLGGNAITACSFALATAGAAVRNEELFLY<br>LARAFHGADKFENLKFRLPTPMVNILNGGKHAGGRLQIQEFMILPKENQPFREKVRCVAE<br>VYQHLGKILAERAGPSAKNVGDEGGFAPNLETADEALNYIEEAIGKAGYKVGEDVFLALDA<br>ASSEFYNSDTKKYEITQQKEFLTSEEMVEYYVQLVNRHPAIISIEDGLEEKDYEGWKLLTE<br>RLGSKIMLVGDDLYTTNTRLIKQGIEEKWANALLLKVNQIGTITEAMNAARMIFNVGQKVIV<br>SHRSGETATTLISDLVVGIGATHIKTGATARGERVSKYNRLLQIEEYLEQHGLLA |

**Supplemental Table S2. X-ray Data collection and processing.** Values for the outer resolution shell are given in parentheses.

| | |
|---|---|
| PDB | 7UGH |
| Ligand | 2-phosphoglycerate (2PG) |
| Diffraction source | APS Beamline 27-ID-F |
| Wavelength (Å) | 0.97872 |
| Temperature (K) | 100 K |
| Detector | Rayonix MX-300 |
| Crystal-detector distance (mm) | 260 |
| Rotation range per image (°) | 1 |
| Total rotation range (°) | 85 |
| Space group | $P4_12_12$ |
| *a*, *b*, *c* (Å) | 96.86, 96.96, 216.64 |
| α, β, γ (°) | 90, 90, 90 |
| Mosaicity (°) | 0.143 |
| Resolution range (Å) | 47.3-1.95 (2.00-1.95) |
| Total No. of reflections | 520,503 (838,442) |
| No. of unique reflections | 75,148 (5432) |
| Completeness (%) | 98.9 (98.5) |
| Redundancy | 6.9 (7.1) |
| $\langle I/\sigma(I)\rangle$ | 19.6 (3.16) |
| $R_{meas}$ | 6.2 (64.6) |
| Overall *B* factor from Wilson plot (Å$^2$) | 32.5 |

**Supplemental Table S3. Crystallographic structure solution and refinement quality statistics**. Values for the outer shell are given in parentheses.

| | |
|---|---|
| PDB | 7UGH |
| Ligand | 2-phosphoglycerate (2PG) |
| Resolution range (Å) | 47.26-1.95 (2.00-1.95) |
| Completeness (%) | 99.0 (98.0) |
| σ cutoff | 1.36 |
| No. of reflections, working set | 75,141 (1956) |
| No. of reflections, test set | 5060 (144) |
| Final $R_{cryst}$ (%) | 15.4 (20.4) |
| Final $R_{free}$ (%) | 19.1 (26.34) |
| No. of non-H atoms | |
| Protein | 6818 |
| Ion | 2 |
| Ligand | 65 |
| Water | 640 |
| Total | 7525 |
| R.m.s. deviations | |
| Bonds (Å) | 0.009 |
| Angles (°) | 0.903 |
| Average *B* factors (Å$^2$) | |
| Protein | 36.9 |
| Ion | 42.6 |
| Ligand | 43.8 |
| Water | 42.7 |
| Ramachandran plot | |
| Most favoured (%) | 98.0 |
| Allowed (%) | 2.0 |

**Supplemental Table S4. Unbiased metabolomics results for amoebae grown in the presence of HEX.** Fold-change (FC) is the mean ratio of peak area of a metabolite detected in the first group divided by the corresponding value in the second group. One-way ANOVA with the Tukey method for post-test was used for metabolite abundance significance test, and the Benjamini and Hochberg method was used for multiple test correction. ND stands for not detected. The metabolite was either not present in the sample or its concentration in sample is smaller than the sensitivity of the analytical platform.

| Name | $t_R$(min) | (-)GLU+GLY vs. GLU | | GLU+HEX vs.GLU | | GLU+GLY+HEX vs. (-)GLU+GLY+HEX | |
|---|---|---|---|---|---|---|---|
| | | FC[a] | *q*-Value[b] | FC[a] | *q*-Value[b] | FC[a] | *q*-Value[b] |
| N-Acetylneuraminic acid | 1.25 | 0.27 | 0.000 | 0.89 | 0.773 | 0.85 | 0.652 |
| 4-Aminobenzoic acid | 1.26 | 3.95 | 0.000 | 1.42 | 0.078 | 0.66 | 0.305 |
| Norethisterone enanthate | 1.89 | 0.05 | 0.000 | 0.95 | 0.868 | 0.94 | 0.921 |
| Uracil | 4.32 | 65.59 | 0.000 | 1.37 | 0.989 | 3.83 | 0.185 |
| 3-Methylhistamine | 4.35 | 2.24 | 0.000 | 1.46 | 0.082 | 0.74 | 0.538 |
| Nicotinic acid | 4.46 | 10.30 | 0.000 | 0.88 | 0.891 | 0.83 | 0.981 |
| Succinate | 4.93 | 1.53 | 0.000 | 0.82 | 0.914 | 0.97 | 0.911 |
| Xanthine | 5.32 | 21.05 | 0.000 | 2.03 | 0.335 | 1.28 | 0.718 |
| Hypoxanthine | 5.34 | 24.87 | 0.000 | 0.64 | 0.666 | 0.51 | 0.818 |
| U (Uridine) | 5.35 | 20.81 | 0.000 | 0.72 | 0.800 | 3.23 | 0.241 |
| Leu-Val | 5.55 | 16.84 | 0.000 | 2.43 | 0.166 | 1.92 | 0.241 |
| 3-(2-Hydroxyethyl)indole | 5.70 | 8.22 | 0.000 | 1.32 | 0.743 | 2.91 | 0.091 |
| 5-Hydroxyindoleacetic acid | 5.90 | 4.42 | 0.000 | 0.44 | 0.043 | 1.15 | 0.902 |
| (S)-(+)-Allantoin | 5.93 | 0.10 | 0.000 | 0.95 | 0.895 | 0.71 | 0.290 |
| Guanine | 6.05 | 10.28 | 0.000 | 0.60 | 0.372 | 1.47 | 0.690 |
| Methionine | 6.26 | 3.10 | 0.000 | 1.47 | 0.055 | 1.15 | 0.936 |
| Creatinine | 6.48 | 0.20 | 0.000 | 1.55 | 0.000 | 0.57 | 0.248 |
| Alanine/Sarcosine | 6.99 | 2.85 | 0.000 | 0.79 | 0.335 | 0.92 | 0.526 |
| S-Adenosylmethionine | 7.82 | 4.14 | 0.000 | 0.97 | 0.178 | 0.87 | 0.267 |
| Aspartic acid | 7.85 | 0.21 | 0.000 | 0.61 | 0.099 | 0.65 | 0.599 |
| Cystathionine | 8.00 | 3.09 | 0.000 | 0.63 | 0.236 | 0.63 | 0.731 |
| Val-arg | 8.55 | 5.33 | 0.000 | 2.12 | 0.302 | 1.06 | 0.921 |
| Arginine | 8.65 | 1.26 | 0.000 | 1.91 | 0.000 | 0.90 | 0.936 |
| Ornithine | 8.73 | 3.04 | 0.000 | 0.86 | 0.493 | 0.98 | 0.725 |
| C4-Carnitine | 16.40 | 0.78 | 0.000 | 0.88 | 0.043 | 0.90 | 0.803 |
| (2,7-Dimethyloctahydro-1H-cyclopenta[c]pyrid | 17.37 | 0.80 | 0.000 | 0.92 | 0.306 | 1.07 | 0.783 |
| trans-4-Hydroxy-L-Proline | 1.08 | 2.77 | 0.009 | 1.17 | 0.323 | 0.73 | 0.690 |
| Glycerophosphocholine | 1.10 | 2.62 | 0.009 | 2.32 | 0.440 | 0.31 | 0.091 |
| Ala-ser | 1.18 | 3.13 | 0.009 | 1.01 | 0.940 | 1.14 | 0.833 |
| Creatine | 1.30 | 0.25 | 0.009 | 1.31 | 0.104 | 0.65 | 0.290 |
| Benzoic acid | 2.38 | 1.69 | 0.009 | 1.06 | 0.335 | 0.94 | 0.902 |
| PS(18:1(9Z)/18:1(9Z)) | 4.67 | 3.66 | 0.009 | 0.68 | 0.138 | 0.72 | 0.536 |
| (2R)-1-[[(2-Aminoethoxy)(hydroxy)phosphoryl]o | 5.13 | 0.50 | 0.009 | 0.68 | 0.138 | 1.07 | 0.867 |
| cAMP (3' 5') | 6.27 | 58.38 | 0.009 | 2.50 | 0.365 | 1.09 | 0.811 |
| 1-Methyladenine | 6.30 | 4.26 | 0.009 | 1.08 | 0.855 | 1.01 | 0.976 |
| Glycyl-L-leucine | 6.37 | 5.35 | 0.009 | 2.01 | 0.082 | 1.21 | 0.921 |
| Tyrosine | 6.65 | 1.81 | 0.009 | 1.26 | 0.333 | 1.03 | 0.731 |
| 5'-S-Methyl-5'-thioadenosine | 10.87 | 3.59 | 0.009 | 1.38 | 0.812 | 1.10 | 0.764 |
| Dehydrowarfarin | 17.79 | 0.82 | 0.009 | 0.92 | 0.587 | 1.01 | 0.994 |
| (5Z)-7,10-Dihydroxy-5-tetradecen-8-ynoic acid | 18.22 | 0.85 | 0.009 | 0.91 | 0.049 | 0.87 | 0.918 |
| 3-[(3-Hydroxyheptanoyl)oxy]-4-(trimethylammo | 18.52 | 0.81 | 0.009 | 0.90 | 0.092 | 0.96 | 0.918 |
| LysoPC(18:3(9Z,12Z,15Z)) | 19.03 | 3.44 | 0.009 | 1.30 | 0.302 | 0.91 | 0.894 |
| 2-(Bromomethyl)-1,3-butadiene | 22.12 | 0.74 | 0.009 | 0.87 | 0.138 | 0.98 | 0.884 |
| FMNH2 | 0.83 | 6.23 | 0.015 | 3.56 | 0.643 | 0.56 | 0.412 |
| 5-GMP | 1.62 | 3.54 | 0.015 | 2.04 | 0.309 | 1.16 | 0.718 |
| Nicotine | 4.85 | 1.59 | 0.015 | 1.19 | 0.840 | 0.74 | 0.731 |
| Pyruvate | 5.37 | 4.96 | 0.015 | 1.14 | 0.910 | 1.19 | 0.674 |
| 11-Aminoundecanoic acid | 5.74 | 2.06 | 0.015 | 1.74 | 0.043 | 0.80 | 0.731 |
| UDP-GAL (UDP-Galactose)/UDP-Glucose | 7.95 | 0.44 | 0.015 | 1.18 | 0.809 | 0.74 | 0.844 |
| 13(S)-HOTrE | 19.60 | 0.71 | 0.015 | 0.91 | 0.614 | 1.27 | 0.808 |
| Valine | 1.63 | 1.62 | 0.019 | 0.98 | 0.910 | 1.05 | 0.887 |
| Nicotinamide | 2.25 | 1.62 | 0.019 | 1.45 | 0.117 | 0.85 | 0.732 |
| Adenine | 5.31 | 3.36 | 0.019 | 1.16 | 0.773 | 1.23 | 0.731 |
| Isoleucine | 6.08 | 1.84 | 0.019 | 1.02 | 0.439 | 1.00 | 0.981 |
| G (Guanosine) | 6.21 | 8.36 | 0.019 | 1.76 | 0.883 | 1.41 | 0.645 |
| C (Cytidine) | 6.69 | 4.13 | 0.019 | 0.97 | 0.992 | 3.96 | 0.091 |
| Adenosine monophosphate | 7.18 | 2.07 | 0.019 | 2.09 | 0.049 | 0.85 | 0.803 |
| P-Hydroxybenzaldehyde | 12.93 | 1.56 | 0.019 | 0.98 | 0.920 | 0.97 | 0.936 |
| 4-nitrocatechol | 14.03 | 2.93 | 0.019 | 0.75 | 0.433 | 1.67 | 0.288 |
| 5-HexN-(tert-Butoxycarbonyl)-L-leucine | 16.56 | 0.85 | 0.019 | 0.97 | 0.643 | 0.97 | 0.783 |
| CMPF | 16.66 | 0.78 | 0.019 | 0.80 | 0.324 | 1.06 | 0.829 |
| 2-[(carboxymethyl)(methyl)amino]-5-methoxyb | 19.58 | 0.74 | 0.019 | 0.85 | 0.323 | 1.06 | 0.441 |
| L-(+)-Valine | 1.14 | 2.65 | 0.022 | 1.81 | 0.035 | 0.66 | 0.091 |
| Hexadecasphinganine | 4.36 | 0.85 | 0.022 | 0.95 | 0.440 | 1.02 | 0.690 |
| Glutamic acid | 7.36 | 0.59 | 0.022 | 0.66 | 0.162 | 0.59 | 0.448 |
| 3-Methyl-1-phenyl-2-butene | 14.89 | 0.69 | 0.022 | 1.08 | 0.940 | 0.93 | 0.918 |
| (+)-Allopumiliotoxin 267A | 18.99 | 1.57 | 0.022 | 1.89 | 0.440 | 0.48 | 0.444 |
| 3-Hydroxyfluorene | 19.07 | 0.83 | 0.022 | 0.91 | 0.055 | 1.04 | 0.936 |
| 1,4-Di-tert-butylbenzene | 19.27 | 0.81 | 0.022 | 0.96 | 0.488 | 1.03 | 0.864 |
| 1-(10Z-heptadecenoyl)-sn-glycero-3-phosphoch | 19.43 | 2.68 | 0.022 | 0.76 | 0.643 | 0.88 | 0.844 |
| Gly-DL-Phe | 6.35 | 2.75 | 0.026 | 0.91 | 0.812 | 1.56 | 0.718 |
| Capryloylglycine | 14.78 | 0.81 | 0.026 | 0.94 | 0.323 | 0.64 | 0.697 |
| 3-[(2,6-Dimethylheptanoyl)oxy]-4-(trimethylam | 19.43 | 0.77 | 0.026 | 0.76 | 0.309 | 1.58 | 0.625 |
| DL-Carnitine | 1.14 | 0.41 | 0.029 | 1.31 | 0.481 | 0.70 | 0.290 |
| L-alpha-Aminoadipic acid | 7.17 | 0.56 | 0.029 | 0.31 | 0.035 | 0.73 | 0.400 |
| (Triethoxymethoxy)ethane | 12.65 | 0.75 | 0.029 | 0.98 | 0.609 | 1.04 | 0.703 |
| 11-Nitro-1-undecene | 17.29 | 0.85 | 0.029 | 0.93 | 0.309 | 1.02 | 0.833 |
| L-Hexanoylcarnitine | 17.30 | 0.73 | 0.029 | 0.97 | 0.870 | 0.88 | 0.560 |
| 2,6-di-tert-butylhydroquinone | 17.87 | 0.76 | 0.029 | 0.84 | 0.241 | 0.99 | 0.837 |
| galegine | 4.47 | 2.15 | 0.033 | 1.47 | 0.312 | 0.80 | 0.625 |
| 1-Palmitoyl-2-linoleoyl PE | 4.68 | 1.70 | 0.033 | 1.27 | 0.633 | 0.82 | 0.567 |
| Glutaric acid | 4.78 | 1.69 | 0.033 | 0.71 | 0.323 | 1.48 | 0.091 |
| Glycerol 3-phosphate | 7.52 | 56.29 | 0.033 | 7.65 | 0.253 | 0.36 | 0.373 |
| Inosine | 5.76 | 15.97 | 0.035 | 1.65 | 0.840 | 2.22 | 0.783 |
| Tetramethyltetrazene | 6.54 | 0.85 | 0.035 | 0.95 | 0.149 | 1.04 | 0.823 |
| Precocene II | 20.09 | 0.85 | 0.035 | 0.94 | 0.346 | 0.96 | 0.809 |
| C17 Sphingosine | 20.40 | 0.74 | 0.035 | 0.93 | 0.391 | 1.03 | 0.811 |
| (2R)-2-Acetoxy-3-(9-decen-1-yloxy)propyl 2-(trir | 5.15 | 0.58 | 0.039 | 0.43 | 0.043 | 0.94 | 0.981 |
| 1-Hydroxy-6,6-dimethyl-3,5,5a,6,9a,9b-hexahy | 18.49 | 0.35 | 0.039 | 0.75 | 0.720 | 2.02 | 0.000 |
| 16-Heptadecynoic acid | 20.16 | 1.16 | 0.039 | 3.65 | 0.570 | 1.45 | 0.645 |
| 2-Phosphoglycerate(2PG)/3-Phosphoglyceric ac | 1.09 | 9.30 | 0.041 | 73.54 | 0.035 | 1.43 | 0.734 |
| N-Acetylputrescine | 6.31 | 6.46 | 0.041 | 1.22 | 0.643 | 1.96 | 0.645 |
| N,N-dimethylarginine | 7.50 | 1.08 | 0.041 | 1.16 | 0.391 | 0.82 | 0.645 |
| 3-[(1E,3E)-1,3-Pentadien-1-yl]-5-phenyl-4,5-dihy | 15.86 | 0.80 | 0.041 | 0.98 | 0.912 | 1.11 | 0.292 |
| 3-methoxyanthranilic acid | 17.95 | 1.43 | 0.041 | 1.03 | 0.323 | 1.32 | 0.645 |
| Arachidonic acid | 1.78 | 3.21 | 0.043 | 1.14 | 0.812 | 1.08 | 0.844 |
| PS(18:0/20:0) | 4.89 | 1.87 | 0.043 | 0.53 | 0.049 | 0.90 | 0.844 |
| DL-N-Acetyltryptophan | 14.75 | 2.00 | 0.043 | 0.52 | 0.209 | 1.07 | 0.811 |
| 2-Aminotetradecanoic acid | 19.20 | 0.81 | 0.043 | 1.01 | 0.982 | 0.97 | 0.811 |
| 2,4-DIMETHYL-2-IMIDAZOLINE | 4.53 | 3.72 | 0.045 | 1.30 | 0.649 | 0.79 | 0.645 |
| C18-Carnitine | 19.14 | 0.88 | 0.045 | 1.36 | 0.934 | 0.82 | 0.766 |
| (9Z,13Z,15Z)-14,18-Dihydroxy-12-oxo-9,13,15- | 19.73 | 0.85 | 0.045 | 0.98 | 0.863 | 1.08 | 0.232 |
| Adipic acid | 10.67 | 0.66 | 0.048 | 0.95 | 0.863 | 1.35 | 0.690 |
| Phenylethyl alcohol | 17.77 | 0.83 | 0.048 | 0.88 | 0.000 | 1.05 | 0.680 |
| A (Adenosine) | 5.32 | 2.98 | 0.052 | 1.33 | 0.633 | 1.43 | 0.000 |
| L-(-)-Asparagine | 7.19 | 1.03 | 0.083 | 1.03 | 0.198 | 0.60 | 0.000 |
| D-Fructose-6-Phosphate(F6P) | 7.84 | 0.63 | 0.124 | 4.26 | 0.043 | 0.77 | 0.947 |
| Dodecenoic acid | 18.12 | 0.82 | 0.178 | 0.92 | 0.049 | 0.82 | 0.055 |
| 3-Methyl-L-histidine | 8.41 | 0.63 | 0.210 | 3.24 | 0.000 | 0.70 | 0.538 |
| Choline | 4.44 | 0.72 | 0.293 | 2.12 | 0.049 | 0.54 | 0.000 |
| Sedoheptulose7-phosphate(SH7P) | 7.94 | 1.37 | 0.297 | 5.84 | 0.000 | 1.80 | 0.833 |
| 1-Oleoyl-2-hydroxy-sn-glycero-3-PE | 4.92 | 1.43 | 0.461 | 0.42 | 0.049 | 1.38 | 0.538 |
| Malate | 1.54 | 0.87 | 0.502 | 0.52 | 0.043 | 1.02 | 0.987 |
| D-Glucose-6-Phosphate(G6P) | 8.07 | 0.79 | 0.627 | 6.83 | 0.043 | 0.90 | 0.997 |
| Mannose 6-phosphate | 1.08 | 1.12 | 0.668 | 8.21 | 0.035 | 0.74 | 0.977 |
| 1D-myo-Inositol 1,2-cyclic phosphate | 7.94 | 0.89 | 0.890 | 5.47 | 0.035 | 1.04 | 0.997 |
| benzoquinone | 7.84 | 0.98 | 0.907 | 5.84 | 0.000 | 0.98 | 0.941 |
| 4-Methyl-2-oxovaleric acid | 12.16 | ND | ND | ND | ND | 2.09 | 0.000 |

**Supplemental Table S5**. **Fold-change in abundance[1] of metabolites in response to carbon source and HEX treatment, relative to growth in glucose alone.** [1]Relative to average intensities of metabolites detected in cells grown under standard (+glc) conditions.

| Metabolite | +HEX | -glc/+gly | -glc/+gly +HEX |
|---|---|---|---|
| Glucose | 0.51 | 0.41 | 0.41 |
| G6P | 7.4 | 0.95 | 13 |
| F6P | 4.5 | 0.83 | 6.5 |
| Gly3P | 17.9 | $1.5 \times 10^2$ | $5.1 \times 10^2$ |
| 2-/3-PG | 78 | 11 | $3.7 \times 10^2$ |
| PYR | 1.2 | 5.8 | 4.3 |

**Supplemental Table S6. Fold-change in abundance of metabolites of cells grown in glycerol after HEX treatment, relative to growth in glycerol alone.** [1]Relative to average intensities of metabolites detected in cells grown in -glc/+gly conditions.

| Metabolite | -glc/+gly +HEX |
|---|---|
| Glucose | 1.2 |
| G6P | 2.9 |
| F6P | 1.7 |
| Gly3P | 26 |
| 2-/3-PG | 7.2 |
| PYR | 0.58 |
